# Effect of autonomous consumption of different sweet substances on the urinary proteome of mice

**DOI:** 10.1101/2024.05.24.595844

**Authors:** Haitong Wang, Youhe Gao

**Affiliations:** Beijing Key Laboratory of Genetically Engineered Drugs and Biotechnology, School of Life Sciences, Beijing Normal University, Beijing, China Beijing 100875)

**Keywords:** urine proteome, sweeteners, sugar metabolism, brain reward pathways

## Abstract

**Objective:** To explore the possible effects on the organism by analysing changes in the urinary proteome of mice after autonomous consumption of different sweet substances.

**Methods:** Urine samples were collected from C57BL/6l mice before and after active consumption of sweet substances. The sweet substances included sucrose, stevia glycosides, acesulfame, and sucralose, which are more widely used worldwide and can elicit a preference response in mice, and the concentrations of the non-nutritive sweeteners were chosen to be those that have been shown by existing studies to have the strongest preference response in mice. The analyses were performed by the non-labelled quantitative proteomics technique of high performance liquid chromatography tandem mass spectrometry (LC-MS/MS), and groups were screened for differential proteins in the urinary proteome for the analysis of protein functions and biological processes; comparisons of urinary proteomes before and after consumption of sweeteners by a single mouse were carried out; and side-by-side comparisons of different sweeteners were made.

**Results and Conclusions:** Urine proteome could reflect the changes in the organism of mice after voluntary consumption of sweeteners, and the effects of different sweeteners on the urine proteome were not the same. Among the four sweeteners, sucralose and sucrose induced the most similar changes in the organism, and steviol glycosides induced the furthest changes in the organism; the changes induced by sucrose, acesulfame, and sucralose were similar, and steviol glycosides induced changes different from those of the other sweeteners. Stevia glycosides induced changes that were different from the other sweeteners. The urinary proteomic proteins that differed among the four sweeteners consumed autonomously by mice included proteins that have been reported to be associated with brain reward circuits, whereas only the urinary proteomic proteins that differed among the four sweeteners consumed autonomously by mice were associated with metabolic processes in large quantities after the voluntary consumption of sucrose, acesulfame, and sucralose, and the urinary proteomic proteins that differed among the four sweeteners consumed autonomously by mice were associated with the assembly of nucleosomes, gene expression, and other processes after the voluntary consumption of stevia glycosides.

## 1 Introduction

Sugar, as a type of carbon hydrate, can be categorised into monosaccharides: glucose, fructose and galactose; and disaccharides: sucrose (Sucrose), lactose and maltose [1]. Sugar did not move into the diet of most people until recent times, becoming an unusual flavouring and providing a large amount of energy and a pleasant sweet taste experience, but human ancestors evolved in a low-sugar environment and had difficulty adapting to the sudden and large addition of sugar to their diet in a short period of time [2 3]. A growing body of research suggests that excessive sugar intake leads to an increased risk of obesity [4], diabetes [5], cardiovascular disease [6], non-alcoholic fatty liver disease [7], dental caries [8], and certain cancers [9]; also, due to the innate sensitivity to sweetness in human beings, which results in the inability to adapt to high levels of sweet stimuli for a short period of time, excessive intake of sugar produces brain reward signals in the brain that go beyond the normal, potentially overriding the self-control mechanisms, leading to addiction, and intense sweet sensory stimuli can exceed the rewards of cocaine stimulation [2 3].

More and more people are aware of the risk of excessive sugar intake, and non-nutritive sweeteners are becoming increasingly popular as sugar substitutes, widely used in the production of products such as food and beverages, due to their no or low calorie content and their ability to provide a more intense sweetness sensory experience [10]. Currently, the most widely used non-nutritive sweeteners worldwide mainly include aspartame, acesulfame, sucralose, stevia glycosides, etc [11]. However, there has been controversy about the safety and efficacy of non-nutritive sweeteners. An 11-year study showed that consumption of two or more servings of beverages containing non-nutritive sweeteners per month increased the risk of coronary heart disease and chronic kidney disease [12 13], and in a study of 17 subjects, an increase in peak plasma glucose concentration and a 20% increase in insulin secretion rate were observed in a glucose tolerance test after the ingestion of sucralose relative to the ingestion of water [14]. In the face of the increasing use of non-nutritive sweeteners, a renewed and effective large-scale evaluation to screen for safe and more popular non-nutritive sweeteners is an urgent issue.

Urine is produced by the filtration of blood through the kidneys for the elimination of metabolic wastes, and is not controlled by the homeostatic regulatory mechanisms of the internal environment, and can be more sensitive to retaining the various small changes produced by the organism[15]. It has been suggested that the search for biomarkers through the urinary metabolome could be used to provide an objective assessment of the intake of non-nutritive sweeteners, thus enhancing population-based studies [16]. However, no studies have been conducted to investigate the effects of sweeteners, including sugar and non-nutritive sweeteners, on the organism through the urine proteome. In contrast to the urinary metabolome, changes in the urinary proteome directly reflect changes in the organism, which, together with its inherent sensitivity, has the potential for direct research on the short- and long-term effects of sugar and non-nutritive sweeteners on the organism. In the present study, the sweetening substances that are currently more widely used worldwide and can cause preference reactions in mice: sucrose, stevia glycosides, acesulfame, sucralose, of which the concentrations of non-nutritive sweeteners are all chosen to be those that have been shown by existing studies to have the strongest preference reactions in mice [17 18], and a comparative study was conducted on the urinary proteome through the collection of urine samples from the mice before and after autonomous consumption of the different sweetening substances. An attempt was made to explore whether the changes in the organism caused by sugar and non-nutritive sweeteners could be reflected in the urine proteome, and then to explore which non-nutritive sweeteners have a lesser effect on the organism and are close to the effect of sucrose, and whether there are any other potential effects of non-nutritive sweeteners on the organism.

## 2 Experimental Methods

### 2.1 Experimental animals

Nine 10-week-old C57BL/6l male mice were purchased from Beijing Viton Lihua Experimental Animal Biotechnology Co. All mice were kept in a standard environment (room temperature (22±1)°C, humidity 65%-70%). All mice were kept in the new environment for three days before starting the experiment, and all experimental operations followed the review and approval of the Ethics Committee of the College of Life Sciences, Beijing Normal University, with the approval number CLS-AWEC-B-2022-003.

### 2.2 Urine sample collection

The sweetening substances selected for this study were sucrose, stevia glycosides, acesulfame, and sucralose, which are more widely used worldwide and can elicit a preference response in mice, and the concentrations of the non-nutritive sweeteners were chosen to be those that have been shown by existing studies to have the strongest preference response in mice [17 18].

#### 2.2.1 Urine sample collection in sucrose group

Four male rats were given water for 1 h after 10 h of water fasting and urine collection for 12 h after water fasting to obtain samples from the sucrose control group; male rats were acclimatised and fed for 2 days; male rats were licked 0.2 g of sucrose powder after 10 h of water fasting and water for 1 h, and urine collection for 12 h after water fasting to obtain samples from the sucrose experimental group, which were temporarily stored in the refrigerator at -80°C.

#### 2.2.2 Collection of urine samples in the stevia glycosides group

Five male rats were given water to collect urine for 12 h after 10 h of water fasting to obtain steviol glycoside control group samples; male rats were acclimatised and fed for 1 day; male rats were given 0.3% steviol glycoside in water to collect urine for 12 h after 10 h of water fasting to obtain steviol glycoside experimental group samples, which were temporarily stored in a -80° refrigerator. Male rats were acclimatised and fed for 2 days.

#### 2.2.3 Urine sample collection in the acesulfame group

Five male rats were given water to collect urine for 12 h after 10 h of water fasting to obtain samples from the acesulfame control group; male rats were acclimatised and fed for 1 day; male rats were given 10 mM acesulfame to collect urine for 12 h after 10 h of water fasting to obtain samples from the acesulfame experimental group, and the samples were stored in the refrigerator at -80°. Male rats were acclimatised and fed for 2 days.

#### 2.2.4 Urine sample collection in sucralose group

Five male rats were given water to collect urine for 12 h after 10 h of water fasting to obtain samples from the sucralose control group; male rats were acclimatised and fed for 1 day; after 10 h of water fasting in male rats, 10 mM sucralose water was given to collect urine for 12 h to obtain samples from the sucralose experimental group, which were temporarily stored in the refrigerator at -80°.

### 2.3 Urine sample processing

Urine protein extraction: Mouse urine samples were removed from the refrigerator at -80 °C and thawed at 4°C. Mouse urine samples were centrifuged at 4°C, 12000×g for 30 min and the supernatants were transferred to new Eppendorf (EP) tubes. Triple volumes of precooled absolute ethanol was added, and the samples were homogeneously mixed and precipitated overnight at −20 °C. The mixture precipitated overnight was centrifuged at 4°C, 12000×g for 30 min, the supernatant was discarded, and the ethanol was volatilized and dried. The precipitated protein was dissolved in lysate buffer (8 mol/L urea, 2 mol/L thiourea, 25 mmol/L dithiothreitol, 50 mmol/L Tris) and centrifuged at 4°C at 12000×g for 30 min, and the supernatant was placed in a new EP tube. The protein concentration was measured by Bradford method.

Enzyme digestion of urine protein: 100 μg urine protein sample was taken into 1.5 mL EP tube, and 25 mmol/L NH4HCO3 solution was added to make the total volume 200 μL. 20 mM Dithiothreitol solution (DTT, Sigma, prepared in 25 mmol/L NH4HCO3 solution) was added and mixed. The metal bath was heated at 37°C for 60 min, and cooled to room temperature. 50 mM Iodoacetamide (IAA, Sigma, prepared in 25 mmol/L NH4HCO3 solution) was added, mixed and reacted at room temperature for 40 min away from light. 200 μL UA solution (8 mol/L urea, 0.1 mol/L Tris-HCl, pH 8.5) was added to the 10 kDa ultrafiltration tube (Pall, Port Washington, NY, USA), Centrifuged at 18°C, 14000×g for 5 min and repeated once; freshly treated samples were added and centrifuged at 18°, 14000×g for 30 min, The urine protein was on the filter membrane. 200 μL UA solution was added, centrifuged at 18°C, 14000×g for 30 min and repeated twice. 25 mmol/L NH4HCO3 solution was added, centrifuged at 18°C, 14000×g for 30 min and repeated twice. Trypsin Gold (Promega, Fitchburg, WI, USA) was added at the ratio of 1:50 for enzyme digestion, and the water bath at 37°C for 15 h. After digestion, the peptides were collected by centrifugation at 4° at 13000×g for 30 min, desalted by HLB solid phase extraction column (Waters, Milford, MA). The peptides were lyophilized with a vacuum dryer, and stored at -20°C.

### 2.4 LC-MS/MS tandem mass spectrometry analyses

The digested samples were reconstituted with 0.1% formic acid, and peptides were quantified with BCA peptide quantification kit. The peptide concentration was then diluted with 0.1% formic acid to 0.5µg/µL. Mixed peptide samples were prepared from 6 µL of each sample and separated by high pH reversed phase peptide separation kit (Thermo Fisher Scientific, Waltham, MA, USA). Ten fractions were collected by centrifugation, lyophilized with a vacuum dryer and reconstituted with 0.1% formic acid. iRT reagent (Biognosys, Switzerland) was added to each fraction and each digested sample with a volume ratio of sample: iRT of 10:1 to calibrate the retention times of the extracted peptide peaks.

For analysis, 1 µg of peptide from each fraction and each digested sample was loaded onto a trap column and separated on a reverse-phase C18 column (50 µm ×150 mm, 2 µm) using the EASY-nLC1200 HPLC system (Thermo Fisher Scientific, Waltham, MA, USA). The elution for the analytical column lasted 90 min with a gradient of 4%–90% mobile phase B (gradient: mobile phase A: 0.1% formic acid, mobile phase B: 80% acetonitrile; flow rate 0.3 µL/min). Peptides were analysed with an Orbitrap Fusion Lumos Tribrid Mass Spectrometer (Thermo Fisher Scientific, Waltham, MA, USA).

To generate the spectrum library, Ten fractions were subjected to mass spectrometry in data-dependent acquisition (DDA) mode, and 10 raw files were generated. Mass spectrometry data were collected in high sensitivity mode. A complete mass spectrometric scan was obtained in the 350–1200 m/z range with a resolution set at 60,000. 10 raw files were imported into Proteome Discoverer software for library construction using Swiss-iRT and Uniprot-Rat databases (version 2.0, Thermo Scientific). In the sucrose group, 36 variable windows Data Independent Acquisition (DIA) methods for DIA mode of each digested sample were set up according to the results of database construction; in the steviol glycosides, acesulfame and sucralose group, 39 variable window Data Independent Acquisition(DIA) methods for DIA mode of each digested sample were set up according to the results of database construction.

Each digested sample was analysed using Data Independent Acquisition (DIA) mode. DIA mode was performed using the DIA method. After every 10 samples, a single DIA analysis of the pooled peptides was performed as a quality control.

### 2.5 Label-free DIA quantitative analysis

Individual sample raw files collected in DIA mode were imported into Spectronaut Pulsar (Biognosys AG, Switzerland) software for analysis. Peptide abundance was calculated by summing the peak areas of the respective fragment ions in MS2. Protein abundance was calculated by summing the respective peptide abundances.

### 2.6 Data analysis

Three technical replicates were performed for each sample and the mean values were taken for statistical analysis.

In this experiment, group analysis was performed to screen for differential proteins by comparing after consumption of sweetening substances with before consumption of sweetening substances. Differential proteins were screened under the following conditions: Fold change (FC) ≥1.5 or ≤0. 67 between groups, and P value <0.05 for two-tailed unpaired t-test analysis. Screened differential proteins were analysed via the Uniprot website (https://www.uniprot.org/) and the DAVID database (https://david.ncifcrf.gov/) for analysis. Relevant proteins involved in brain reward circuits were summarised by searching the literature related to brain reward circuits in the Pubmed database (https://pubmed.ncbi.nlm.nih.gov) among the differential proteins analysed in cohorts to retrieve proteins related to brain reward circuits that have been reported.

Meanwhile, this experiment was conducted to screen differential proteins by comparative analysis between single mice after and before consumption of sweeteners, and the differential proteins were screened under the following conditions: multiplicity of change (FC, Fold change) ≥ 1.5 or ≤ 0.67 between groups, and P-value < 0.05; and shared differential proteins common to all the five samples were counted. The screened shared differential proteins were analysed via the Uniprot website (https://www.uniprot.org/) and retrieved in single mouse urine proteomic differential proteins that have been reported to be associated with brain reward circuits.

## 3 Experimental results and discussion

### 3.1 Analysis of the urinary protein composition group in the sucrose group

#### 3.1.1 Differential proteins

The sucrose experimental group was compared with the sucrose control group urine proteome, screening differential protein conditions were: FC ≥1.5 or ≤0.67, P<0.05. The results showed that 65 differential proteins could be identified in the sucrose experimental group compared with the sucrose control group, and the differential proteins were ranked in the order of FC from the smallest to the largest, and searched by Uniprot, the results were as shown in Table 1.

**Table 1.**
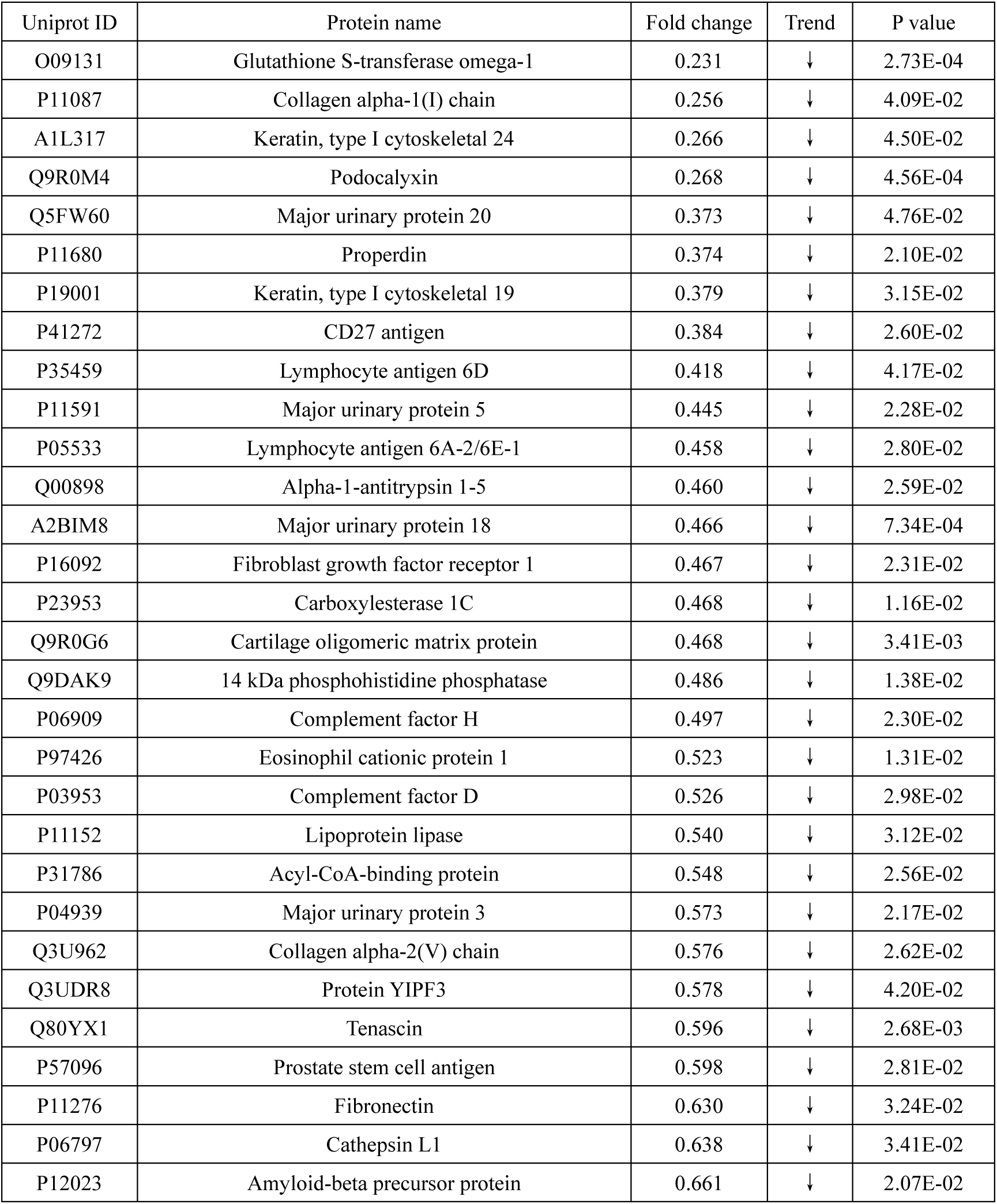
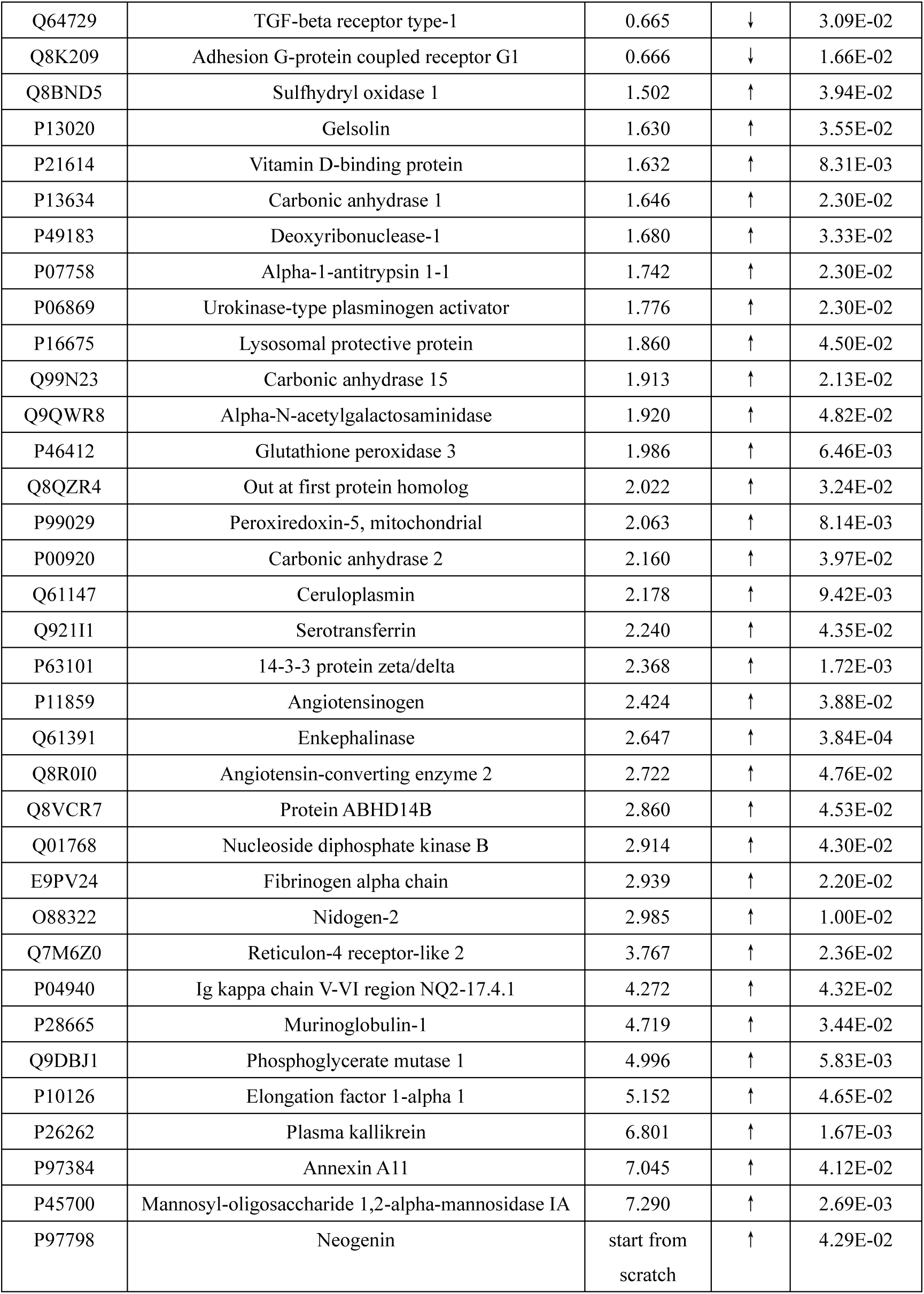
Differential proteins in the urine proteome of mice in the sucrose experimental group and sucrose control group.

#### 3.1.2 Analysis of differential protein function

The 65 identified differential proteins were analysed by DAVID database for molecular functions and biological processes, and the results are shown in Figure 1.

**Fig. 1.**
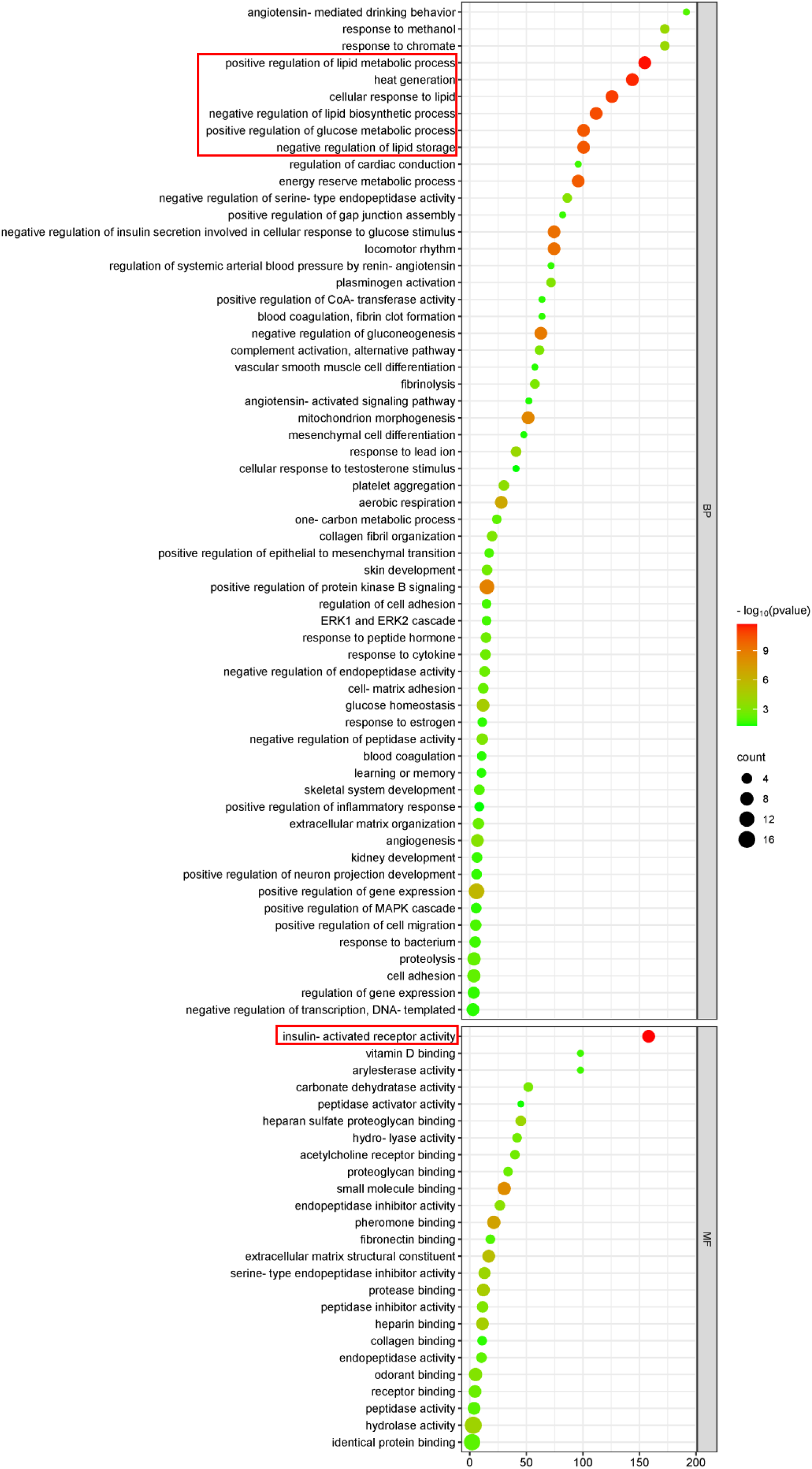
Analysis of molecular functions and biological processes of differential proteins in sucrose group mice

In order of p-value from smallest to largest, 12 of the 18 biological processes ranked in the top 30% showed changes brought about by sugar and lipid metabolism and energy production, such as: positive regulation of lipid metabolism processes, heat production, cellular response to lipids, negative regulation of lipid biosynthesis processes, positive regulation of glucose metabolism processes, negative regulation of lipid storage, and energy reserve metabolism processes, Negative regulation of insulin secretion involved in cellular response to glucose stimuli, negative regulation of gluconeogenesis, mitochondrial morphogenesis, aerobic respiration, glucose homeostasis, etc.; in descending order of p-value, the first molecular function of which is insulin receptor activation, and the rest of the molecular functions are also mainly related to molecular binding, activation of various types of enzymes, and odour binding. This is likely to be related to the metabolic response resulting from sucrose consumption. It is noteworthy, however, that the differential proteins were equally enriched for neuronal projection development for positive regulation, learning or memory, and other neurally relevant biological processes, which may be related to the preference response to sweet taste sensation.

#### 3.1.3 Differential proteins and brain reward circuits

To further investigate whether the urinary proteome differential proteins have brain reward circuit-related proteins, the functions and biological processes involved in sucrose group differential proteins were searched by Uniprot to find whether they are related to the key proteins that have been reported to be involved in brain reward circuits [19], and the results are shown in Table 2.

**Table 2.** Urinary proteomic differential proteins and brain reward circuits in sucrose group mice.

| The Brain's Reward Circuitry |  |
| --- | --- |
| Glutamate | √ |
| Dopamine |  |
| Gamma-aminobutyric acid / GABA | √ |
| Substance P | √ |
| Enkephalin | √ |

Mouse urine proteome difference proteins were retrieved with proteins associated with brain reward circuits.

Acyl-CoA-binding protein is able to replace the recognition site of benzodiazepine located on γ-aminobutyric acid type A receptor of the neurosurgical drug diazepam, and can act as a neuropeptide to regulate the action of γ-aminobutyric acid receptor; it is involved in biological processes such as learning memory, positive regulation of synaptic transmission, and proliferation of glial cells.

Procathepsin L catalyses the processing of the hormone proenkephalin into the active enkephalin neurotransmitter in secretory vesicles of neuroendocrine chromaffinophilic cells; involved in enkephalin processing, neurodevelopment and other biological processes.

Amyloid-beta precursor protein acts as a cell surface receptor on the surface of neurons to perform physiological functions related to neurite growth, neurite adhesion, and axonogenesis; it is involved in biological processes such as learning and memory, glutamate receptor signalling pathways, associative learning, and neuronal differentiation.

Carbonic anhydrase 2 is involved in biological processes such as the positive regulation of γ-aminobutyric acidergic synaptic transmission.

Angiotensinogen is involved in biological processes such as associative learning and negative regulation of neuronal apoptotic processes.

14-3-3 protein zeta/delta is expressed in regions such as glutamatergic synapses and is involved in biological processes such as the regulation of synaptic maturation and recognition of synaptic targets.

Enkephalinase has the ability to cleave opioid peptides such as Met-enkephalin and Leu-enkephalin, and also catalyses the cleavage of bradykinin, Substance P, and neurotensin peptides; it is involved in biological processes such as learning and memory, catabolism of Substance P, and positive regulation of neurodevelopment.

### 3.2 Individual analyses of the pre-sucrose urinary proteome after consumption of sucrose by individuals in the sucrose group revealed a total of one brain reward circuit-associated protein

The urinary proteome of four mice after sucrose consumption was compared with the urinary proteome of before sucrose consumption, respectively, and the condition of screening for differential proteins was FC ≥1.5 or ≤0.67, and the t-test of P<0.05 was used to count differential proteins shared by the four mice, and the results are shown in Figure 2.

**Fig. 2.**
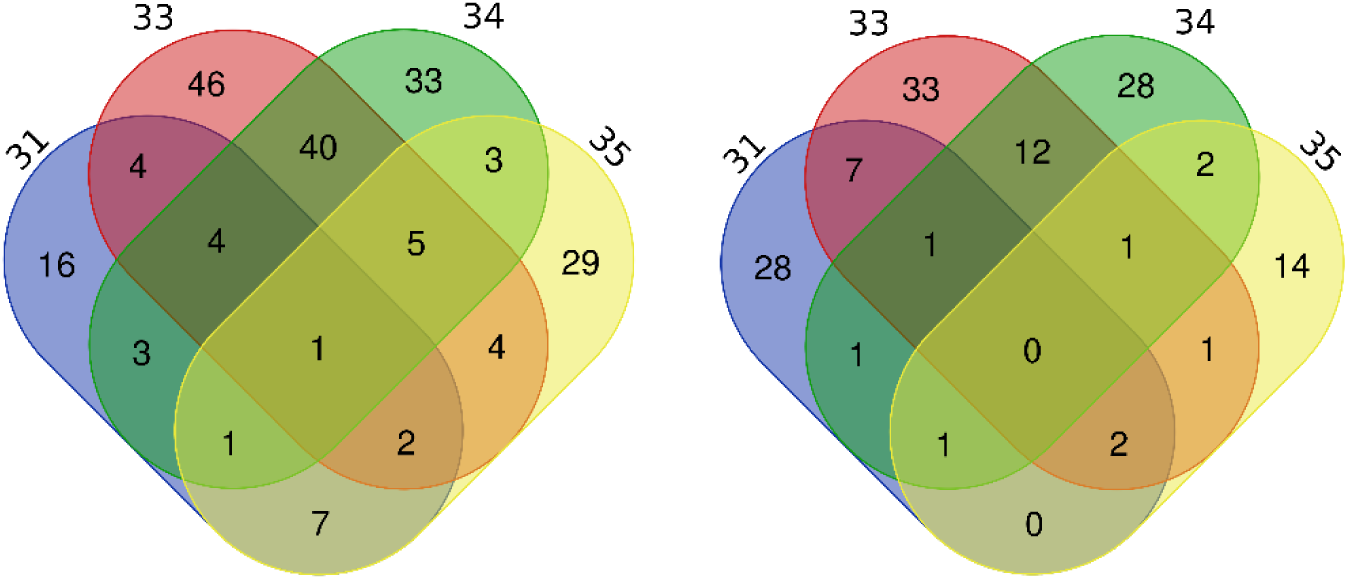
Total up-regulated differential proteins (left) down-regulated differential proteins (right) in front of individuals in the sucrose group after sucrose consumption

There was a total of 1 up-regulated differential protein in 4 mice. Functions of the proteins and biological processes involved were retrieved by Uniprot. Enkephalinase is associated with brain reward circuits and has the ability to cleave opioid peptides such as; Met-enkephalin and Leu-enkephalin, and also catalyses the cleavage of Bradykinin, Substance P, and Neurotensin peptides; and is involved in learning memory, Substance P catabolism, positive regulation of neurodevelopment and other biological processes.

The urine proteomic differential proteins of individual mice before and after sucrose consumption were also searched for brain reward circuit-related proteins that were present in the individual mouse urine proteomic differential proteins, as shown in Table 3.

**Table 3.**
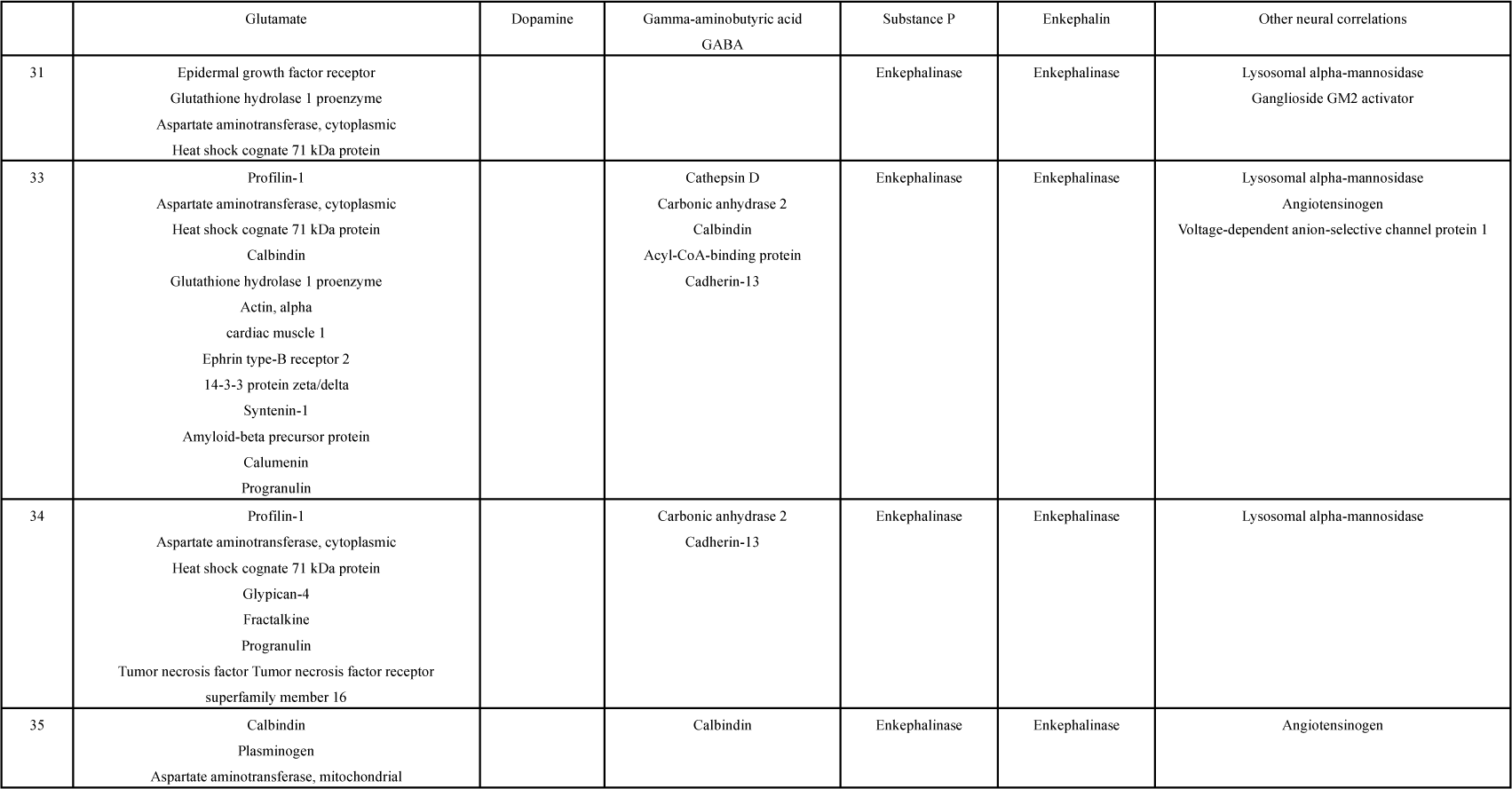
Proteins associated with brain reward circuits in the proteome difference proteins of the anterior urine proteins of individuals in the sucrose group after sucrose consumption.

### 3.3 Compositional analysis of urinary proteins in the stevia glycosides group

#### 3.3.1 Differential proteins

The urine proteome of the stevia glycosides experimental group was compared with that of the stevia glycosides control group, and the conditions for screening differential proteins were FC ≥1.5 or ≤0.67, with P<0.05. The results showed that 66 differential proteins could be identified in the stevia glycosides experimental group compared with that of the stevia glycosides control group, and the differential proteins were arranged in the order of FC from the smallest to the largest, and retrieved through the Uniprot for searching, and the results are shown in Table 4.

**Table 4.**
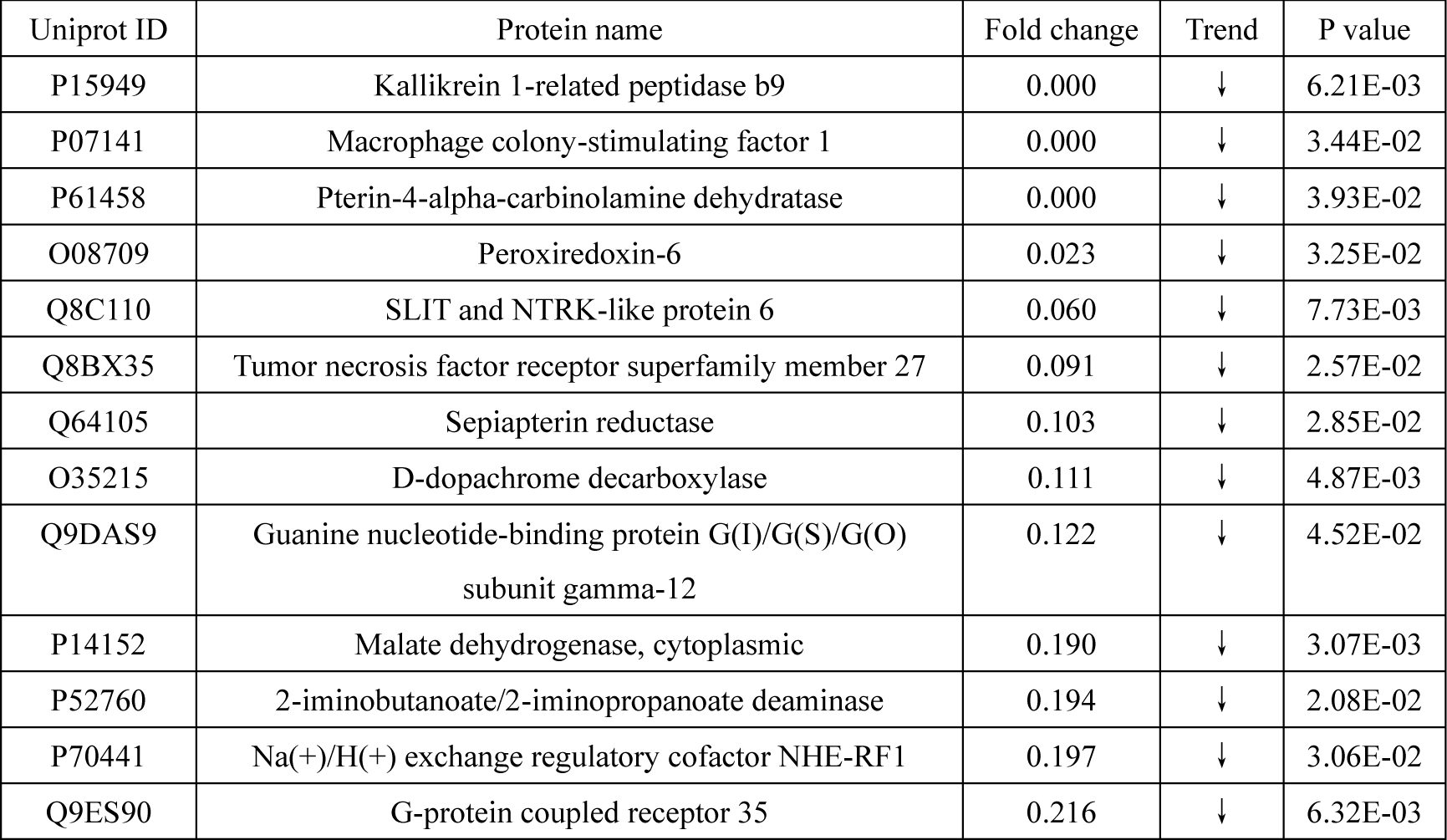
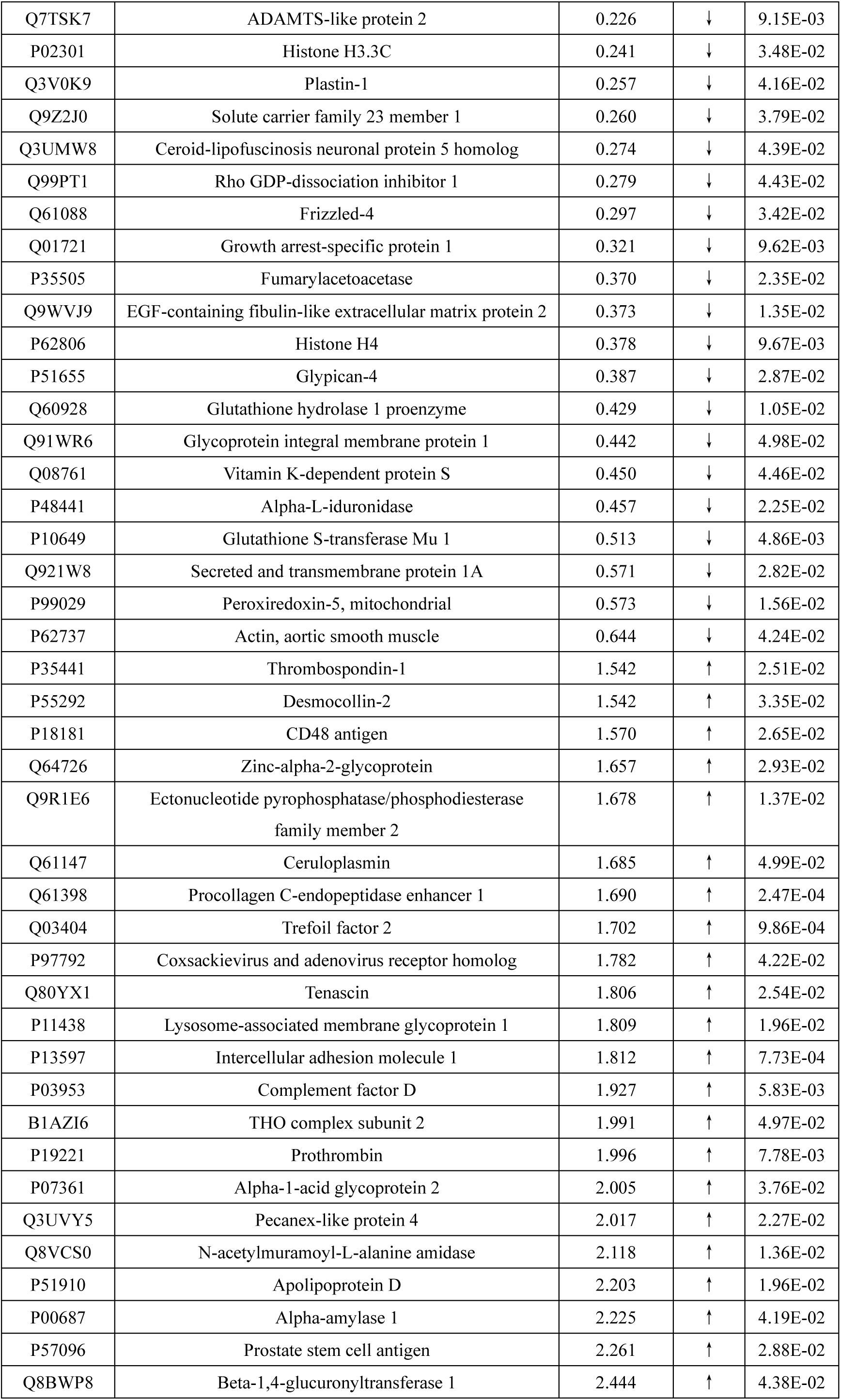
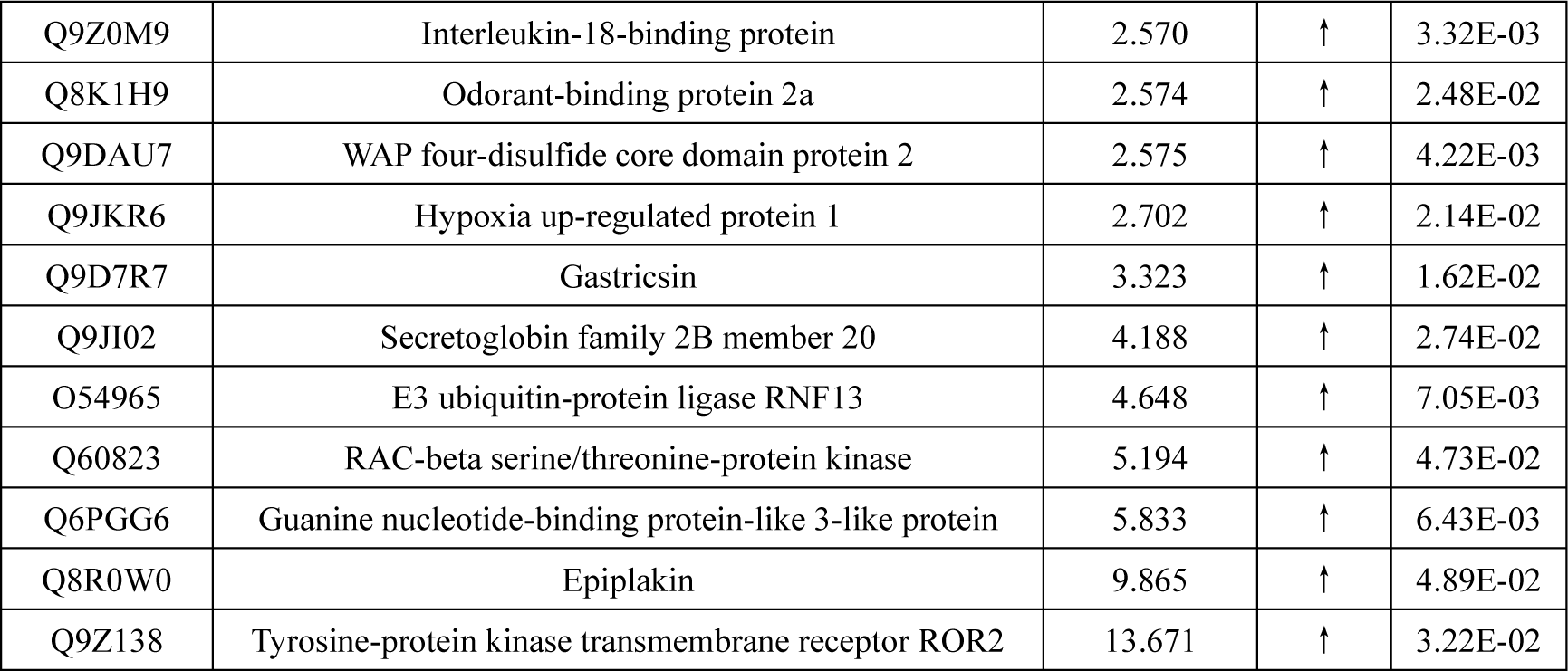
Differential proteins in the urine proteome of mice in the stevia glycosides experimental group and the stevia glycosides control group.

#### 3.3.2 Analysis of differential protein function

The 66 identified differential proteins were analysed by DAVID database for molecular functions and biological processes, and the results are shown in Figure 3.

**Figure 3.**
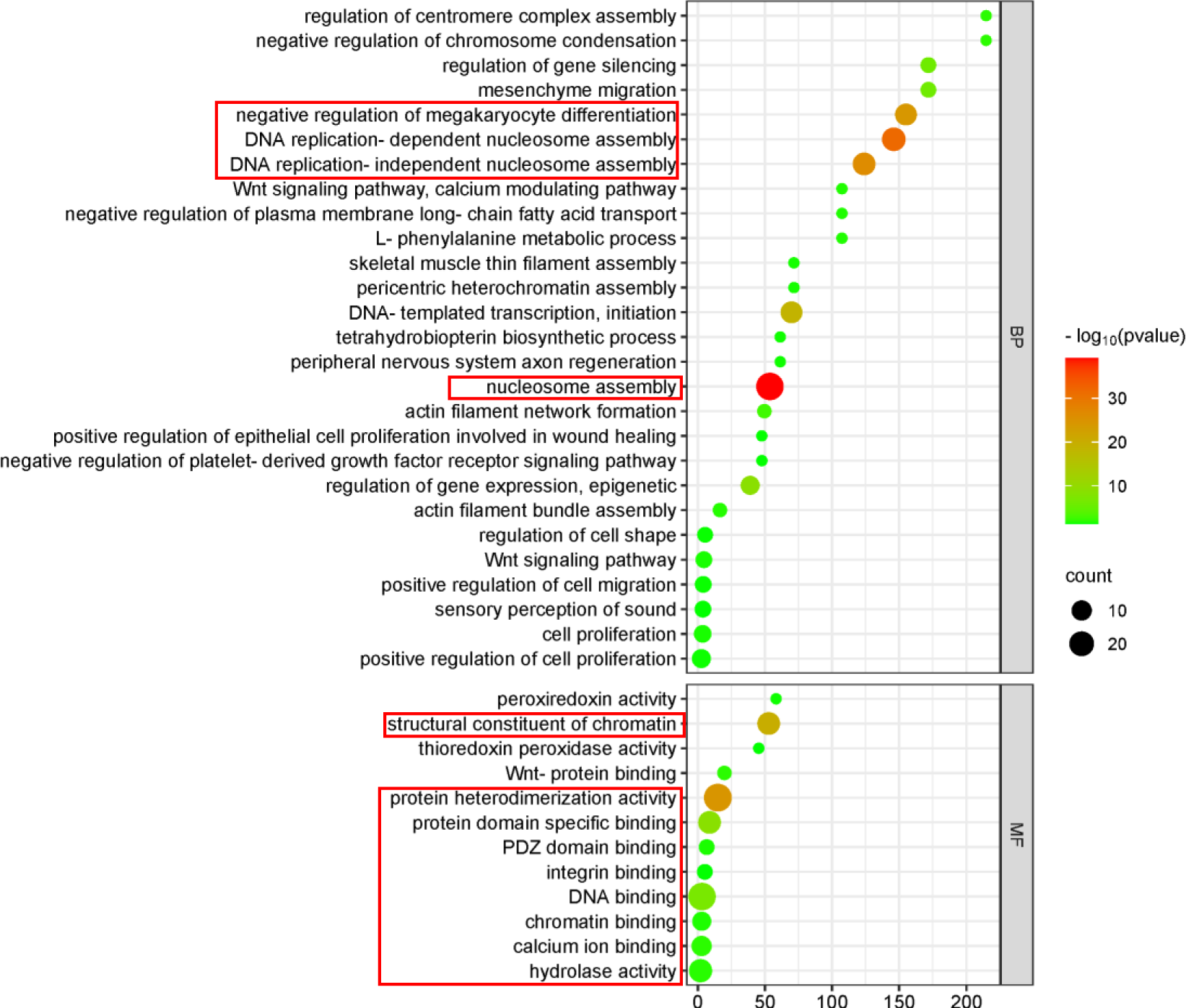
Analysis of molecular functions and biological processes of differential proteins in stevia glycosides group mice

In order of p-value from smallest to largest, 11 of the 13 biological processes in the top 50% of the list showed changes brought about by nucleosome assembly, gene expression, and cell division, such as: nucleosome assembly, DNA replication-dependent nucleosome assembly, DNA replication-independent nucleosome assembly, transcription of DNA templates, initiation, regulation of gene expression, epigenetics, regulation of gene silencing, actin filament network formation, negative regulation of chromosome cohesion, regulation of filament complex assembly, actin filament bundle assembly, etc., Unlike the sucrose group, which exhibits a large number of metabolism-related biological processes, this is the first time that nucleosome-related changes in urinary proteome differential proteins have been seen in mice after autonomous consumption of stevia glycosides, and they are not found in any other sweeteners; sorted by p-value from smallest to largest, in which the top four molecular functions were protein heterodimerisation activity, structural components of chromatin, protein structural domain-specific binding, and DNA binding, and the molecular functions were also mainly related to DNA and protein activities; insulin-related molecular functions, which were significantly present in the sucrose group, were not shown in the stevia glycosides group. Notably, however, the differential proteins were similarly enriched for axon regeneration in the peripheral nervous system, a neurally relevant biological process, which may be related to the preference response to sweet taste perception.

#### 3.3.3 Differential proteins and brain reward circuits

To further investigate whether the urinary proteome differential proteins have brain reward circuit-associated proteins, the functions and biological processes involved in the steviol glycoside group differential proteins were searched by Uniprot to find out whether they are related to the key proteins that have been reported to be associated with brain reward circuits, and the results are shown in Table 5.

**Table 5.** Urinary proteome-differentiated proteins and brain reward circuits in stevia glycosides group mice.

| The Brain's Reward Circuitry |  |
| --- | --- |
| Glutamate | √ |
| Dopamine | √ |
| Gamma-aminobutyric acid / GABA | √ |
| Substance P |  |
| Enkephalin |  |

Mouse urine proteome difference proteins were retrieved with proteins associated with brain reward circuits.

Sepiapterin reductase is involved in biological processes such as cell morphogenesis and dopamine metabolism during neuronal differentiation.

Na(+)/H(+) exchange regulatory cofactor NHE-RF1 functions in dopamine receptor binding and γ-aminobutyric acid transmembrane transport, and is involved in biological processes such as adenylate cyclase-activated dopamine receptor signalling pathway and phospholipase c-activated dopamine receptor signalling pathway.

Frizzled-4 is expressed in glutamatergic synapses and is involved in biological processes such as positive regulation of dendritic morphogenesis and positive regulation of neuronal projection dendriticity.

Glypican-4 is expressed at glutamatergic synapses, presynaptic membranes, and other locations, and is involved in the regulation of postsynaptic specialised membranes by neurotransmitter receptor localisation, presynaptic assembly regulation, synaptic membrane adhesion, and other biological processes.

Glutathione hydrolase 1 proenzyme has glutathione hydrolase activity and is involved in the catabolism of glutathione, the metabolic process of glutamate and the response to alcohol.

Tyrosine-protein kinase transmembrane receptor ROR2 is expressed in dendrites, glutamatergic synapses, neuronal cytosol, and other locations, and is involved in astrocyte development, regulation of chemical synaptic transmission, positive regulation of glutamatergic synaptic transmission, positive regulation of neuronal projection development, and regulation of postsynaptic organisation, among other biological Processes.

### 3.4 Individual analyses of the proteome of the anterior urine of individuals in the stevia glycosides group after stevia glycosides consumption

The urine proteomes of the five mice after steviol glycosides consumption were compared with the urine proteomes before steviol glycosides consumption, respectively, and the conditions for screening differential proteins were FC ≥1.5 or ≤0.67, and the P<0.05 was used to statistically count differential proteins shared by the five mice, and the results are shown in Figure 4.

**Figure 4.**
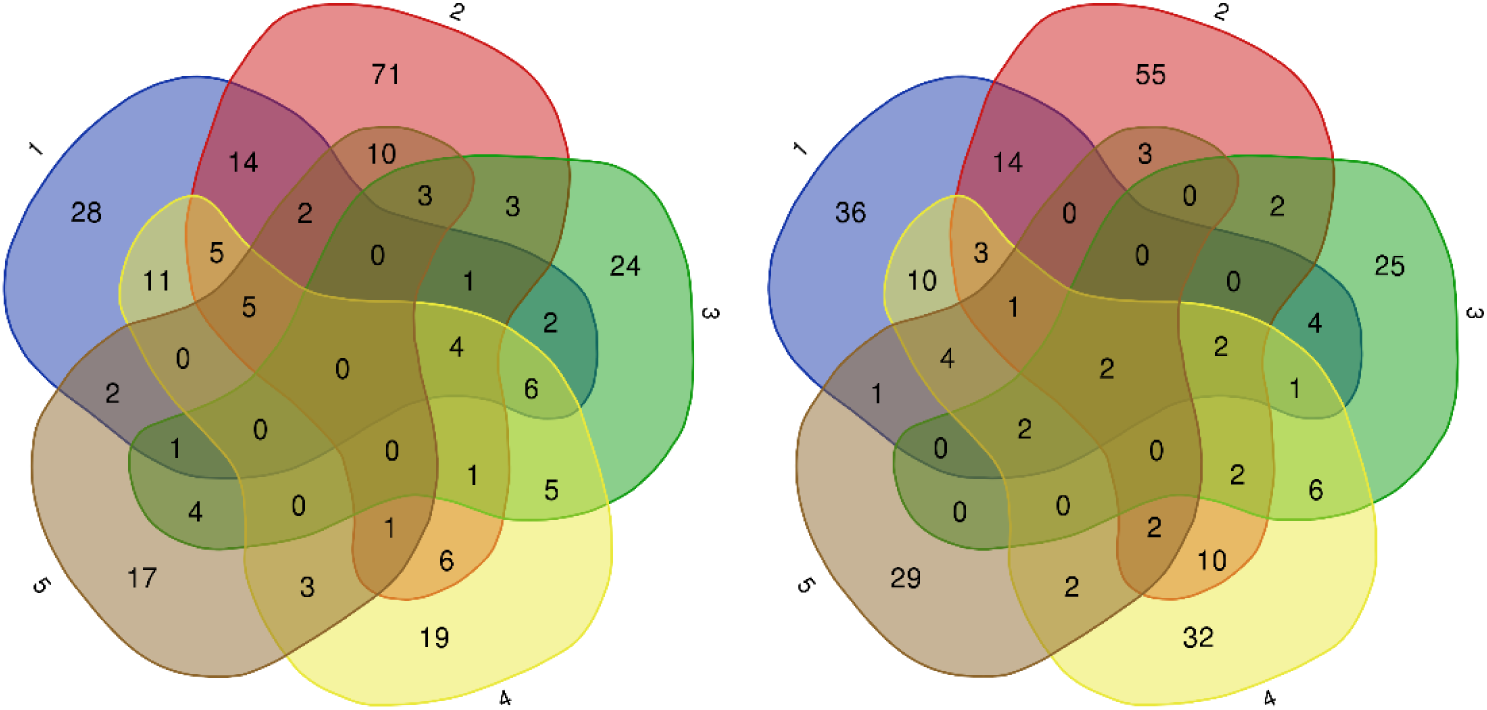
Pre-shared up-regulated differential proteins (left) down-regulated differential proteins (right) after stevia glycosides consumption by individuals in the stevia glycosides group

A total of two down-regulated differential proteins, 2-iminobutanoate/2-iminopropanoate deaminase and Glutathione hydrolase 1 proenzyme, were identified in five mice.Retrieving the function of the proteins and the biological processes involved by Uniprot, the 2-iminobutanoate/2-iminopropanoate deaminase has the function of catalysing the hydrolysis of enamine/imine intermediates formed during normal metabolism deaminating the enamine/imine intermediates, facilitating the recruitment of ribonuclease P/MRP complex to promote ribonucleic acid endocytosis, and is involved in mRNA catabolism, negative translational regulation, and other biological processes; Glutathione hydrolase 1 proenzyme has glutathione hydrolase activity and is involved in glutathione catabolism, glutamate metabolic processes, and response to alcohol.

The urine proteomic differential proteins of individual mice before and after steviol glycoside consumption were also searched for brain reward circuit-related proteins that were present in the individual mouse urine proteomic differential proteins, as shown in Table 6.

**Table 6.**
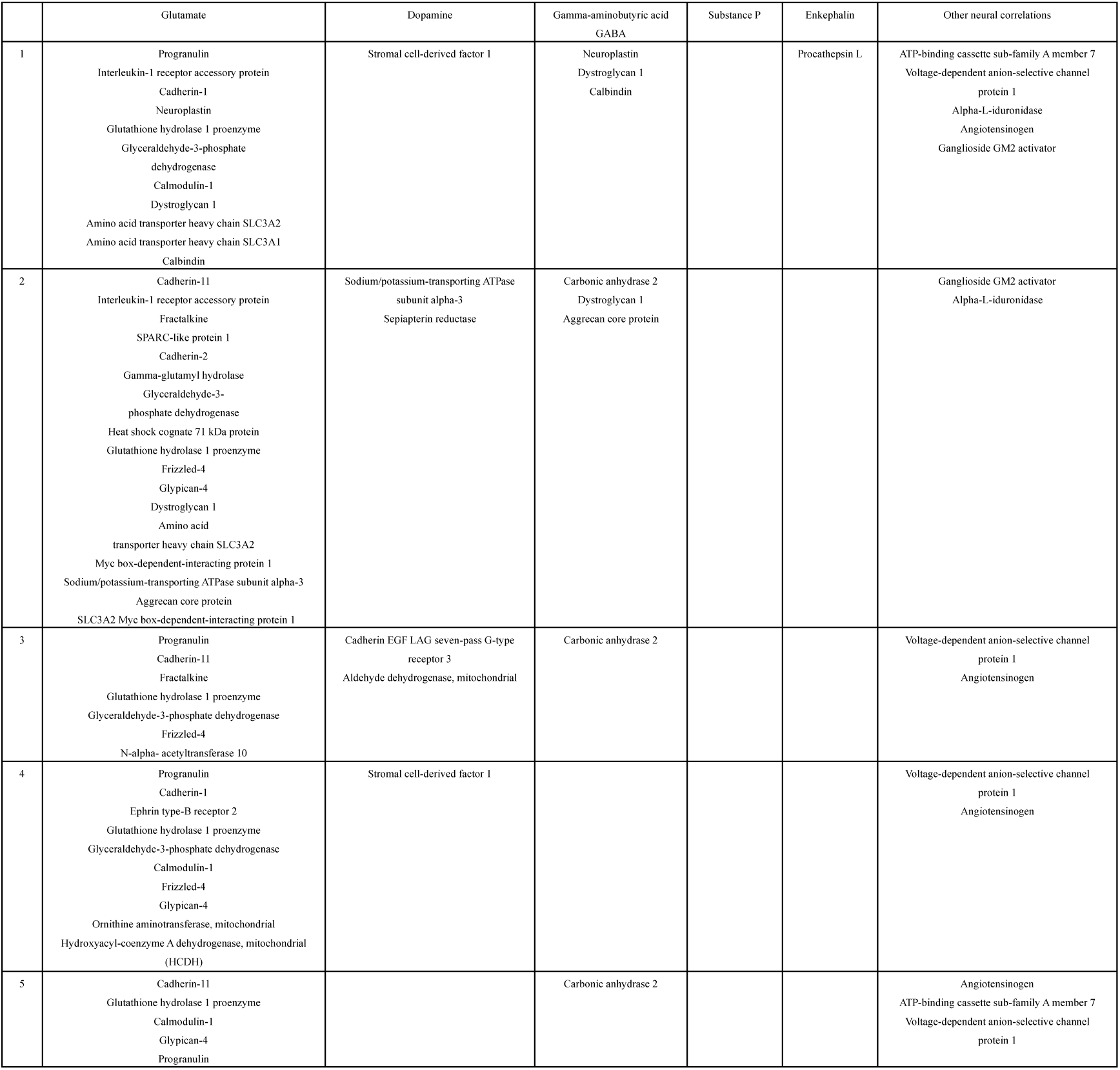
Proteins associated with brain reward circuits in the proteomic differences in urine proteins of individuals in the stevia glycosides group before and after stevia glycosides consumption.

### 3.5 Compositional group analysis of urinary proteins in the acesulfame group

#### 3.5.1 Unsupervised clustering analysis of the total urine proteome of the Acesulfame group

Unsupervised cluster analysis of the total urine proteome of the acesulfame experimental group and the acesulfame control group was performed, and the results are shown in Figure 5. The total proteome can be distinguished before and after the consumption of acesulfame in mice, and the urine proteome of mice changed significantly after the consumption of acesulfame.

**Fig. 5.**
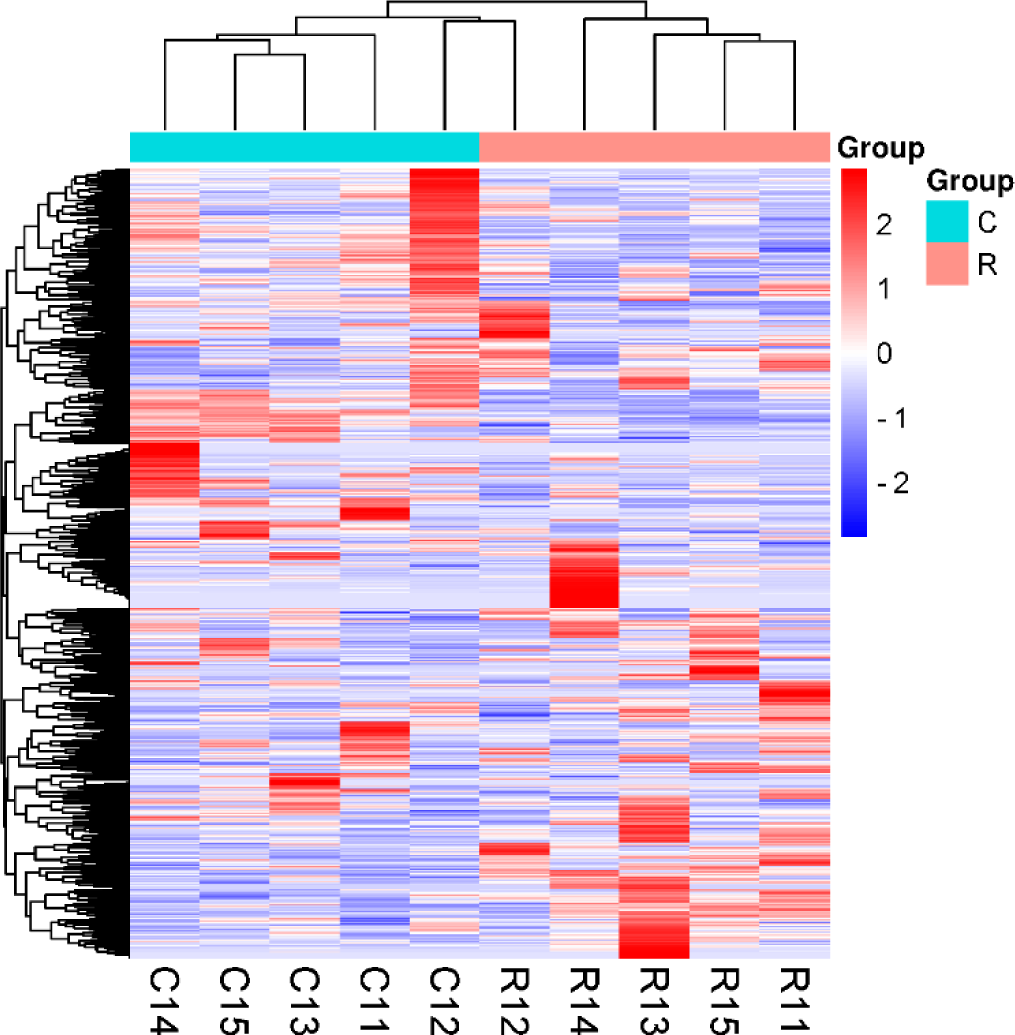
Unsupervised clustering of the total urine proteome of the acesulfame group

#### 3.5.2 Differential proteins

The Acesulfame experimental group was compared with the urine protein of Acesulfame control group, and the conditions for screening differential proteins were: FC ≥1.5 or ≤0.67, P<0.05. The results showed that 93 differential proteins could be identified in the Acesulfame experimental group as compared with the Acesulfame control group, and the differential proteins were ranked in the order of FC from the smallest to the largest, and searched by Uniprot, which was used to search for differential proteins. The results are shown in Table 7.

**Table 7.** Differential proteins in urine proteome of mice from acesulfame experimental group and acesulfame control group.

| Uniprot ID | Protein name | Fold change | Trend | P value |
| --- | --- | --- | --- | --- |
| O35215 | D-dopachrome decarboxylase | 0.00 | ↓ | 7.84E-04 |
| Q61247 | Alpha-2-antiplasmin | 0.00 | ↓ | 1.58E-02 |
| Q14BI7 | ATP-dependent RNA helicase TDRD9 | 0.00 | ↓ | 2.31E-02 |
| P55849 | Desmocollin-1 | 0.00 | ↓ | 2.50E-02 |
| Q8VEM8 | Phosphate carrier protein, mitochondrial | 0.00 | ↓ | 2.64E-02 |
| P35802 | Neuronal membrane glycoprotein M6-a | 0.00 | ↓ | 4.52E-02 |
| P52760 | 2-iminobutanoate/2-iminopropanoate deaminase | 0.01 | ↓ | 1.50E-02 |
| P61458 | Pterin-4-α-carbinolamine dehydratase | 0.02 | ↓ | 2.37E-06 |
| Q9DAS9 | Guanine nucleotide-binding protein G(I)/G(S)/G(O) subunit gamma-12 | 0.03 | ↓ | 9.92E-03 |
| O09051 | Guanylate cyclase activator 2B | 0.08 | ↓ | 2.79E-02 |
| P99029 | Peroxiredoxin-5, mitochondrial | 0.09 | ↓ | 1.17E-03 |
| P70441 | Na(+)/H(+) exchange regulatory cofactor<br>NHE-RF1 | 0.10 | ↓ | 1.74E-02 |
| P35505 | Fumarylacetoacetase | 0.12 | ↓ | 3.36E-03 |
| Q9JLT2 | Trehalase | 0.15 | ↓ | 2.72E-02 |
| P26043 | Radixin | 0.19 | ↓ | 2.57E-02 |
| Q7M6Z0 | Reticulon-4 receptor-like 2 | 0.22 | ↓ | 2.35E-02 |
| P12658 | Calbindin | 0.22 | ↓ | 4.78E-02 |
| Q8K426 | Resistin-like gamma | 0.25 | ↓ | 1.35E-02 |
| Q60928 | Glutathione hydrolase 1 proenzyme | 0.25 | ↓ | 1.24E-02 |
| P81117 | Nucleobindin-2 | 0.26 | ↓ | 1.18E-02 |
| Q08761 | Vitamin K-dependent protein S | 0.30 | ↓ | 5.15E-03 |
| Q91XE4 | N-acyl-aromatic-L-amino acid<br>amidohydrolase (carboxylate-forming) | 0.30 | ↓ | 1.43E-02 |
| Q9ES90 | G-protein coupled receptor 35 | 0.32 | ↓ | 7.29E-03 |
| Q64449 | C-type mannose receptor 2 | 0.38 | ↓ | 8.50E-03 |
| P10649 | Glutathione S-transferase Mu 1 | 0.39 | ↓ | 3.48E-02 |
| Q9QXD6 | Fructose-1,6-bisphosphatase 1 | 0.39 | ↓ | 4.16E-02 |
| O09133 | Submaxillary gland androgen-regulated<br>protein 2, isoform alpha | 0.42 | ↓ | 3.62E-02 |
| Q60932 | Voltage-dependent anion-selective channel<br>protein 1 | 0.43 | ↓ | 1.69E-03 |
| Q6RKD8 | Leucine-rich repeat transmembrane protein<br>FLRT1 | 0.45 | ↓ | 3.82E-03 |
| Q91WV7 | Neutral and basic amino acid transport protein<br>rBAT | 0.49 | ↓ | 3.12E-02 |
| O08992 | Syntenin-1 | 0.49 | ↓ | 3.30E-03 |
| Q9CQ60 | 6-phosphogluconolactonase | 0.50 | ↓ | 3.36E-02 |
| Q8VC60 | Beta-galactosidase-1-like protein | 0.50 | ↓ | 3.82E-03 |
| Q9JHY4 | WAP four-disulfide core domain protein 15B | 0.51 | ↓ | 7.56E-04 |
| Q91ZI0 | Cadherin EGF LAG seven-pass G-type<br>receptor 3 | 0.52 | ↓ | 5.31E-04 |
| P70699 | Lysosomal alpha-glucosidase | 0.54 | ↓ | 6.08E-05 |
| O89051 | Integral membrane protein 2B | 0.54 | ↓ | 2.11E-02 |
| Q02819 | Nucleobindin-1 | 0.55 | ↓ | 5.13E-03 |
| P48441 | Alpha-L-iduronidase | 0.55 | ↓ | 4.30E-02 |
| Q9CZT5 | Vasorin | 0.55 | ↓ | 3.47E-02 |
| P11152 | Lipoprotein lipase | 0.57 | ↓ | 9.54E-03 |
| Q8C050 | Ribosomal protein S6 kinase alpha-5 | 0.57 | ↓ | 2.18E-02 |
| Q9QWR8 | Alpha-N-acetylgalactosaminidase | 0.58 | ↓ | 1.14E-03 |
| O35298 | Acyloxyacyl hydrolase | 0.60 | ↓ | 7.23E-03 |
| P17563 | Methanethiol oxidase | 0.63 | ↓ | 1.31E-02 |
| P23780 | Beta-galactosidase | 0.63 | ↓ | 1.48E-03 |
| O35448 | Lysosomal thioesterase PPT2 | 0.63 | ↓ | 1.34E-03 |
| Q91Y97 | Fructose-bisphosphate aldolase B | 0.65 | ↓ | 1.80E-02 |
| Q9DC11 | Plexin domain-containing protein 2 | 1.54 | ↑ | 4.80E-02 |
| Q91V08 | C-type lectin domain family 2 member D | 1.58 | ↑ | 1.85E-02 |
| P01027 | Complement C3 | 1.60 | ↑ | 2.12E-03 |
| Q61398 | Procollagen C-endopeptidase enhancer 1 | 1.61 | ↑ | 3.20E-03 |
| Q9R1E6 | Ectonucleotide<br>pyrophosphatase/phosphodiesterase family<br>member 2 | 1.67 | ↑ | 1.91E-02 |
| Q80YX1 | Tenascin | 1.68 | ↑ | 4.06E-02 |
| Q61147 | Ceruloplasmin | 1.72 | ↑ | 3.04E-03 |
| P14094 | Sodium/potassium-transporting ATPase<br>subunit beta-1 | 1.72 | ↑ | 3.17E-03 |
| O35305 | Tumor necrosis factor receptor superfamily<br>member 11A | 1.84 | ↑ | 4.66E-02 |
| P28665 | Murinoglobulin-1 | 1.89 | ↑ | 1.76E-02 |
| Q9ESD1 | Prostasin | 1.89 | ↑ | 1.78E-02 |
| P54763 | Ephrin type-B receptor 2 | 1.92 | ↑ | 4.38E-02 |
| Q3UVY5 | Pecanex-like protein 4 | 1.95 | ↑ | 2.08E-02 |
| P18572 | Basigin | 1.96 | ↑ | 4.27E-02 |
| P31809 | Carcinoembryonic antigen-related cell<br>adhesion molecule 1 | 1.99 | ↑ | 1.17E-02 |
| P51910 | Apolipoprotein D | 2.01 | ↑ | 3.56E-03 |
| P20029 | Endoplasmic reticulum chaperone BiP | 2.04 | ↑ | 3.30E-02 |
| P20108 | Thioredoxin-dependent peroxide reductase,<br>mitochondrial | 2.06 | ↑ | 3.94E-02 |
| P13597 | Intercellular adhesion molecule 1 | 2.08 | ↑ | 7.88E-04 |
| Q05909 | Receptor-type tyrosine-protein phosphatase<br>gamma | 2.11 | ↑ | 4.39E-03 |
| P61110 | Kidney androgen-regulated protein | 2.14 | ↑ | 1.85E-02 |
| Q9CQW3 | Vitamin K-dependent protein Z | 2.21 | ↑ | 6.07E-03 |
| P35846 | Folate receptor alpha | 2.22 | ↑ | 4.92E-02 |
| P08249 | Malate dehydrogenase, mitochondrial | 2.24 | ↑ | 1.01E-02 |
| Q9D8B7 | Junctional adhesion molecule C | 2.28 | ↑ | 2.81E-02 |
| P48614 | Protein Wnt-10b | 2.29 | ↑ | 2.92E-02 |
| P57096 | Prostate stem cell antigen | 2.69 | ↑ | 9.27E-03 |
| Q00724 | Retinol-binding protein 4 | 2.85 | ↑ | 3.94E-02 |
| Q61495 | Desmoglein-1-alpha | 2.94 | ↑ | 2.13E-02 |
| Q08423 | Trefoil factor 1 | 3.15 | ↑ | 3.49E-02 |
| P26618 | Platelet-derived growth factor receptor alpha | 3.32 | ↑ | 4.83E-02 |
| P35461 | Lymphocyte antigen 6G (Fragment) | 3.75 | ↑ | 2.64E-02 |
| P04441 | H-2 class II histocompatibility antigen<br>gamma chain | 4.28 | ↑ | 4.45E-02 |
| Q9R007 | C-type lectin domain family 5 member A | 4.39 | ↑ | 6.03E-06 |
| Q64729 | TGF-beta receptor type-1 | 4.55 | ↑ | 4.63E-02 |
| Q9DAU7 | WAP four-disulfide core domain protein 2 | 4.59 | ↑ | 1.78E-02 |
| Q811B3 | A disintegrin and metalloproteinase with thrombospondin motifs 12 | 4.61 | ↑ | 3.82E-02 |
| P16092 | Fibroblast growth factor receptor 1 | 5.44 | ↑ | 4.02E-02 |
| P31725 | Protein S100-A9 | 5.53 | ↑ | 1.83E-02 |
| Q60823 | RAC-beta serine/threonine-protein kinase | 5.83 | ↑ | 3.06E-02 |
| Q9CR33 | MANSC domain-containing protein 1 | 6.10 | ↑ | 1.68E-02 |
| O54965 | E3 ubiquitin-protein ligase RNF13 | 10.79 | ↑ | 4.26E-02 |
| Q9JM99 | Proteoglycan 4 | 16.79 | ↑ | 3.09E-02 |
| P51885 | Lumican | 25.22 | ↑ | 3.02E-03 |
| P07901 | Heat shock protein HSP 90-alpha | 30.83 | ↑ | 3.49E-02 |

#### 3.5.3 Differential protein function analysis

The 93 identified differential proteins were analysed by DAVID database for molecular functions and biological processes, and the results are shown in Figure 6.

**Fig. 6.**
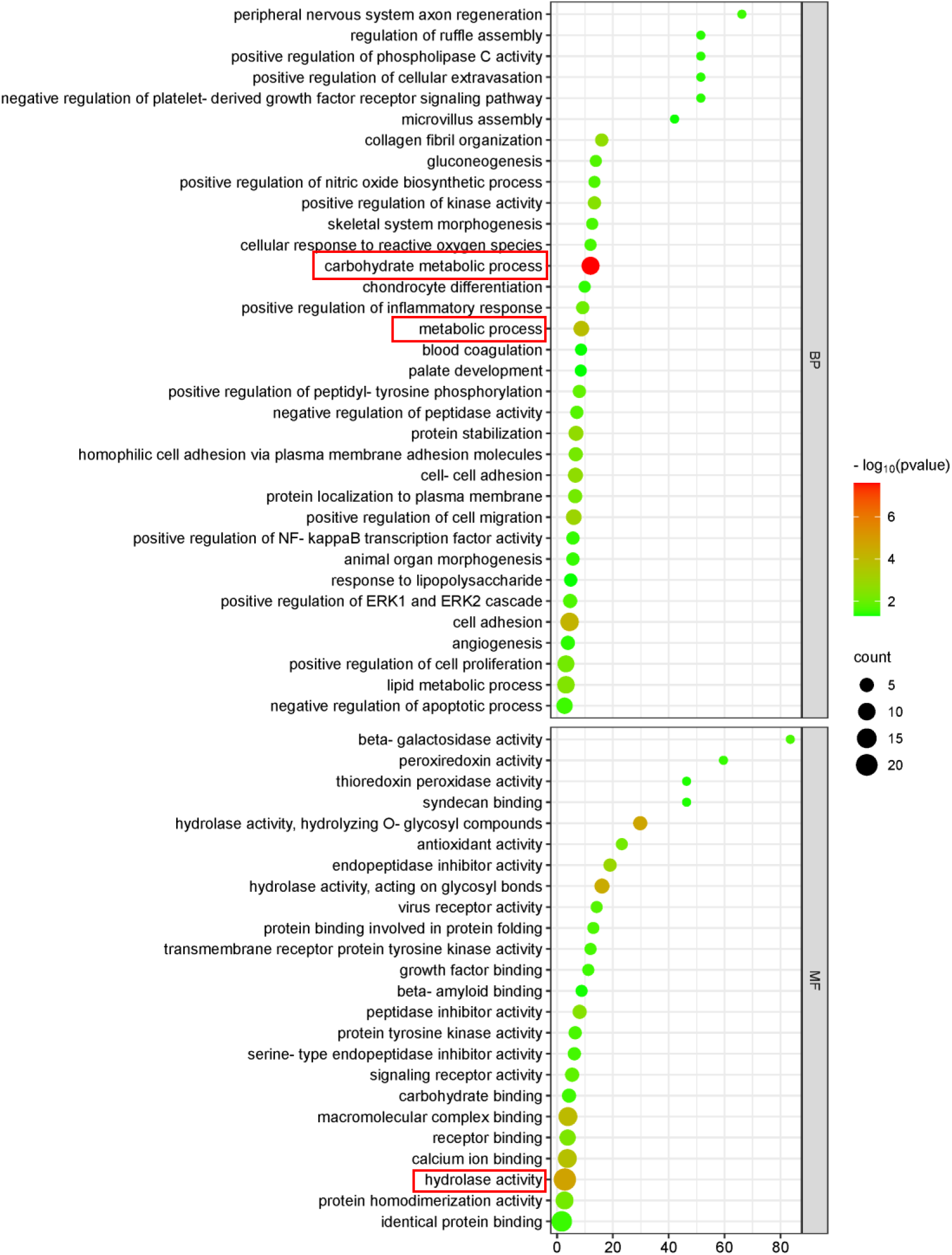
Analysis of molecular functions and biological processes of differential proteins in mice of the acesulfame group

In order of p-value from smallest to largest, among the top 30% of the 10 biological processes, 3 biological processes were involved in sugar and lipid metabolism, such as: carbohydrate metabolism process, lipid metabolism process, metabolism process, etc., which was a smaller proportion than that of the sucrose group; in order of p-value from smallest to largest, among them, the third ranked molecular function was related to the activity of hydrolytic enzymes, especially glycosyl bond hydrolase, which was significantly appeared in the sucrose group Insulin-related molecular functions were not shown in the acesulfame group. Notably, the differential proteins were equally enriched for neural-related biological processes such as axonal regeneration in the peripheral nervous system, which may be related to the preference response to sweet taste sensation.

#### 3.5.4 Differential proteins and brain reward circuits

To further investigate whether the urinary proteome differential proteins have proteins related to brain reward circuits, the functions and biological processes involved in the anserine group differential proteins were searched by Uniprot to find whether they are related to the key proteins that have been reported to be involved in brain reward circuits, and the results are shown in Table 8.

**Table 8.** Urinary proteomic differential proteins and brain reward circuits in mice of the acesulfame group.

| The Brain's Reward Circuitry |  |
| --- | --- |
| Glutamate | √ |
| Dopamine | √ |
| Gamma-aminobutyric acid / GABA | √ |
| Substance P |  |
| enkephalin |  |

Mouse urine proteome difference proteins were retrieved with proteins associated with brain reward circuits.

Neuronal membrane glycoprotein M6-a plays a role in neuronal plasticity and is involved in neural differentiation, including the differentiation and migration of neural stem cells, in the growth of neurons and filopodia, in filopodial motility, and possibly in synapse formation. It is expressed in regions such as axonal growth cones, dendritic spines, glutamatergic synapses, neuronal cytosol, parallel fibre and Purkinje cell synapses, and presynaptic active zone membranes, and is involved in biological processes such as neuronal migration, neuronal projection development, neuronal projection morphogenesis, regulation of synaptic organisation, and synaptic assembly.

Na(+)/H(+) exchange regulatory cofactor NHE-RF1 functions in dopamine receptor binding and γ-aminobutyric acid transmembrane transport, and is involved in biological processes such as adenylate cyclase-activated dopamine receptor signalling pathway and phospholipase c-activated dopamine receptor signalling pathway.

Calbindin is expressed in axons, dendrites, dendritic spines, gamma-aminobutyric acidergic synapses, glutamatergic synapses, neuronal cytosol, and other regions, and is involved in biological processes such as the regulation of synaptic plasticity, the regulation of pre and post-synaptic cytoplasmic calcium ion concentration through calcium binding, the regulation of pre-synaptic cytoplasmic calcium ion concentration, long term memory, short term memory, and so on.

Glutathione hydrolase 1 proenzyme has glutathione hydrolase activity and is involved in the catabolism of glutathione, the metabolic process of glutamate and the response to alcohol.

Amino acid transporter heavy chain SLC3A1 is involved in the transmembrane transport of glutamate.

Syntenin-1 is involved in biological processes such as ionophilic glutamate receptor binding and presynaptic assembly.

Cadherin EGF LAG seven-pass G-type receptor 3 is a receptor that plays an important role in intercellular signalling during the formation of the nervous system; it is involved in biological processes such as axonal fasciculation, axon guidance of dopaminergic neurons, motor neuron migration, involvement of planar cell polarity pathways in axon guidance, and axon guidance of 5-hydroxytryptophan neurons. Involved in.

Ephrin type-B receptor 2 is expressed in regions of axons, dendrites, dendritic spines, glutamatergic synapses, neuronal cytosol, post-synaptic membranes, pre-synaptic membranes, and synapses, and has the function of regulating dendritic spine development and maturation and stimulating excitatory synapse formation. Involved in axonogenesis, CNS projection neuron axonogenesis, joint neuron axon guidance, dendritic spine morphogenesis, learning and memory, neuronal projection contraction, positive regulation of dendritic spine morphogenesis, positive regulation of long term neuronal synaptic plasticity, positive regulation of long term synaptic enhancement, positive regulation of synaptic assembly, positive regulation of synaptic plasticity, post-synaptic membrane assembly, modulation of axonogenesis, neural synaptic plasticity regulation, regulation of synaptic assembly, and the role of the trans-synaptic complex in the regulation of synaptic transmission in biological processes.

### 3.6 Individual analyses of the proteome of the anterior urine of individuals in the acesulfame group after consumption of acesulfame

The urine proteomes of the five mice after consuming acesulfame were compared with those before consuming acesulfame, respectively, and the conditions for screening differential proteins were FC ≥1.5 or ≤0.67, and the P<0.05 was used to count differential proteins shared by the five mice, and the results are shown in Figure 7.

**Fig. 7.**
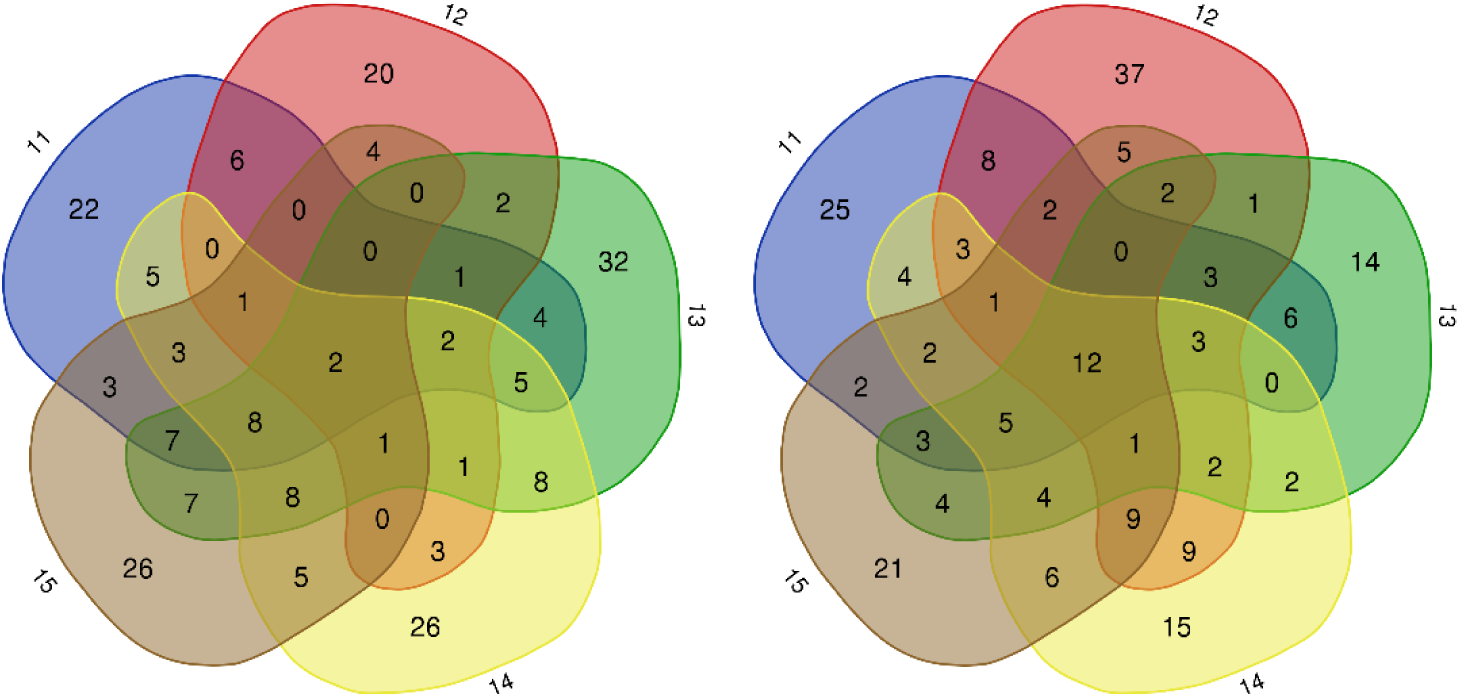
Total up-regulated differential proteins (left) down-regulated differential proteins (right) in front after consumption of acesulfame by individuals in the acesulfame group

There were a total of 2 up-regulated differential proteins and 12 down-regulated differential proteins in 5 mice, as shown in Table 7.

**Table 7.**
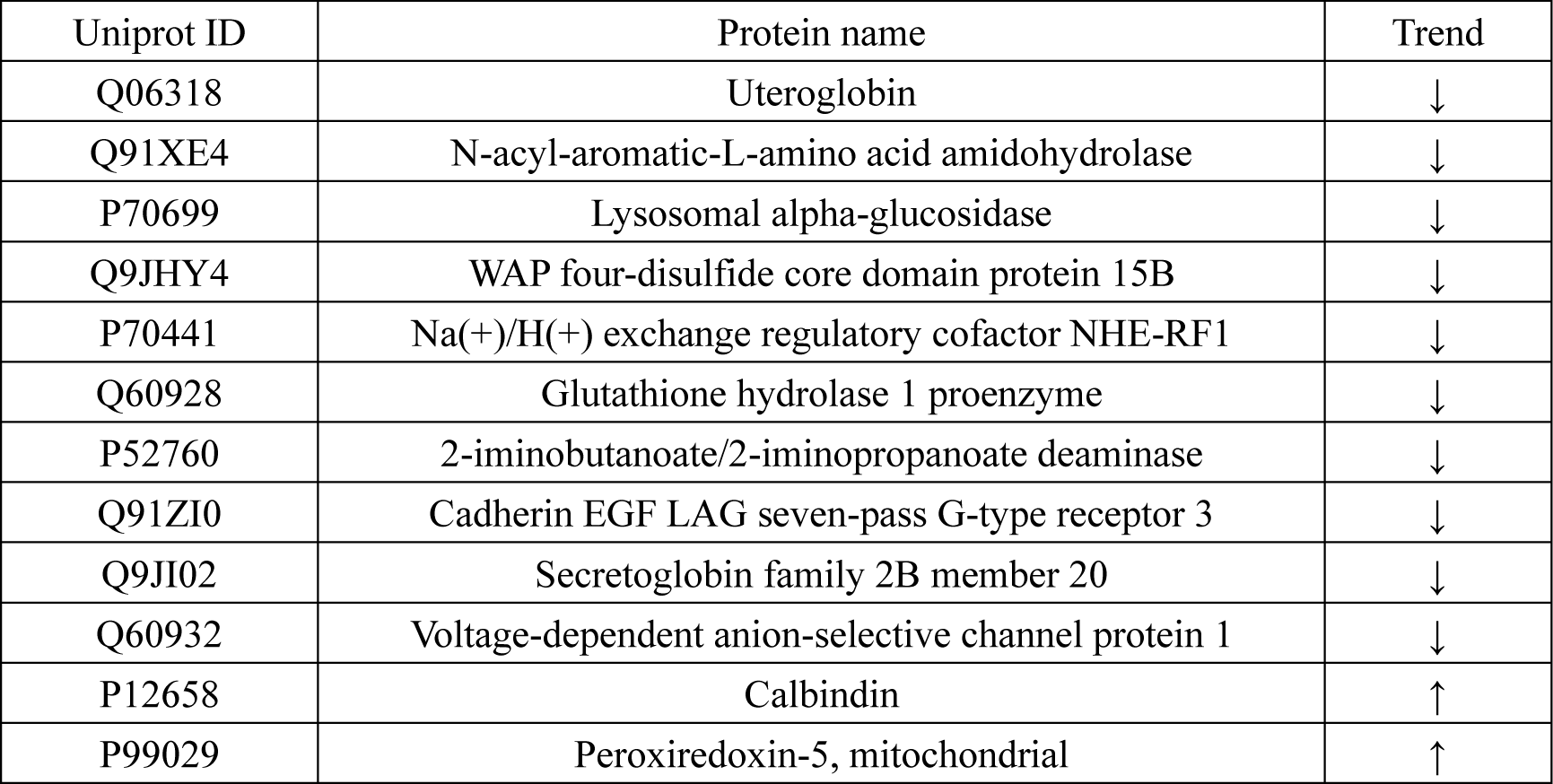
Pre-shared differential proteins after consumption of acesulfame by individuals in the acesulfame group.

The functions and biological processes of the proteins were retrieved by Uniprot, among which Na(+)/H(+) exchange regulatory cofactor NHE-RF1 has the functions of dopamine receptor binding and γ-aminobutyric acid transmembrane transporter, and is involved in the biological processes such as adenylyl cyclase-activated dopamine receptor signalling pathway, phospholipase c-activated dopamine receptor signalling pathway; Cadherin EGF LAG seven-pass G-type receptor 3 is a receptor that plays an important role in intercellular signaling during the formation of the nervous system; and is involved in the axonal signaling process. Cadherin EGF LAG seven-pass G-type receptor 3 is a receptor that plays an important role in intercellular signalling during the formation of the nervous system; it is involved in axonal fasciculation, dopaminergic neuron axon guidance, motor neuron migration, planar cell polarity pathway axon guidance, and 5-hydroxytryptamine neuron axon guidance. Calbindin is expressed in axons, dendrites, dendritic spines, gamma-aminobutyric acid synapses, glutamatergic synapses, and neuronal cytosol, and is involved in the regulation of synaptic plasticity, the regulation of pre and post-synaptic cytoplasmic calcium concentrations through calcium binding, the regulation of pre-synaptic cytoplasmic calcium concentration, long-term memory, and short-term memory. memory, and other biological processes. This may be related to the sweet taste preference response and brain reward circuits.Glutathione hydrolase 1 proenzyme has glutathione hydrolase activity and is involved in the catabolism of glutathione, the metabolism of glutamate and the response to alcohol.

The urine proteomic differential proteins of individual mice before and after consuming acesulfame were also searched for brain reward circuit-related proteins that were present in the individual mouse urine proteomic differential proteins, as shown in Table 9.

**Table 9.**
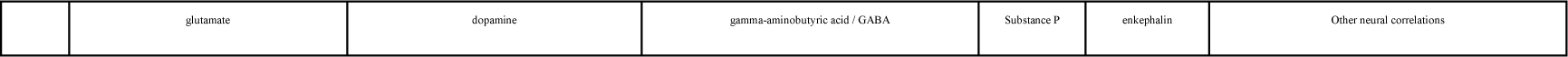
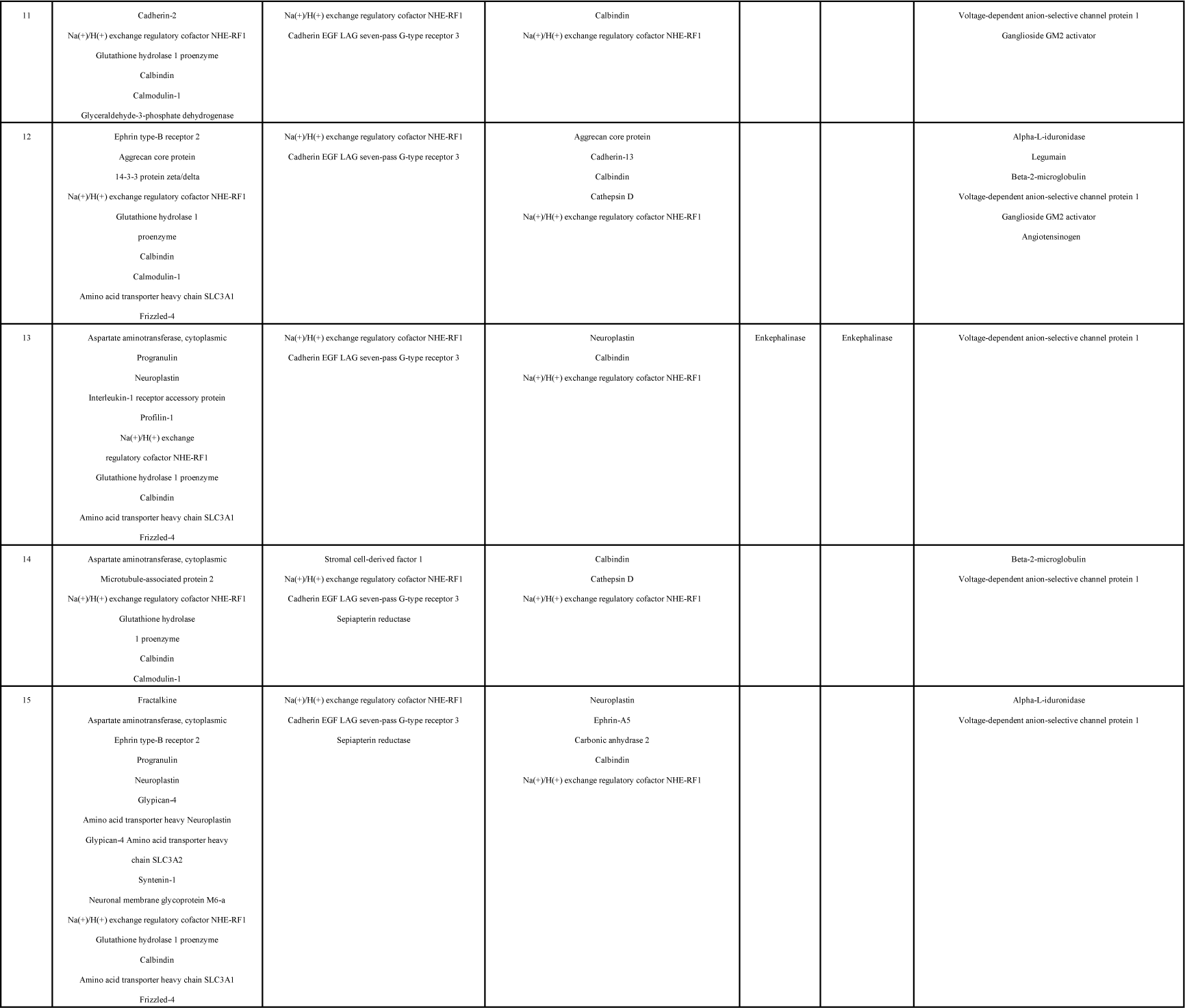
Proteins associated with brain reward circuits in proteomic differences in urine proteins before and after consumption of acesulfame in individuals of the acesulfame group.

### 3.7 Component group analysis of urinary proteins in the sucralose group

#### 3.7.1 Differential proteins

Sucralose experimental group was compared with the sucralose control group for urine protein, and the conditions for screening differential proteins were: FC ≥1.5 or ≤0.67, t-test P<0.05. The results showed that 83 differential proteins could be identified in the sucralose experimental group compared with the sucralose control group, and the differential proteins were arranged in the order of FC from smallest to the largest, and retrieved by Uniprot The search was performed and the results are shown in Table 10.

**Table 10.**
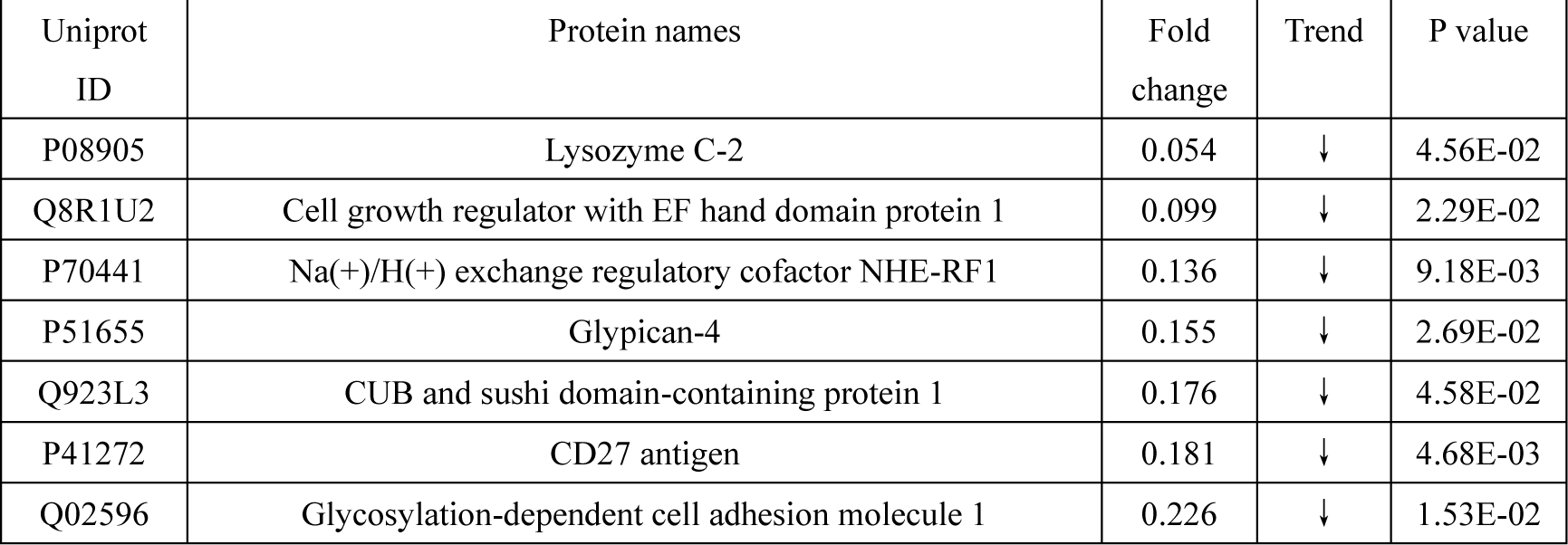
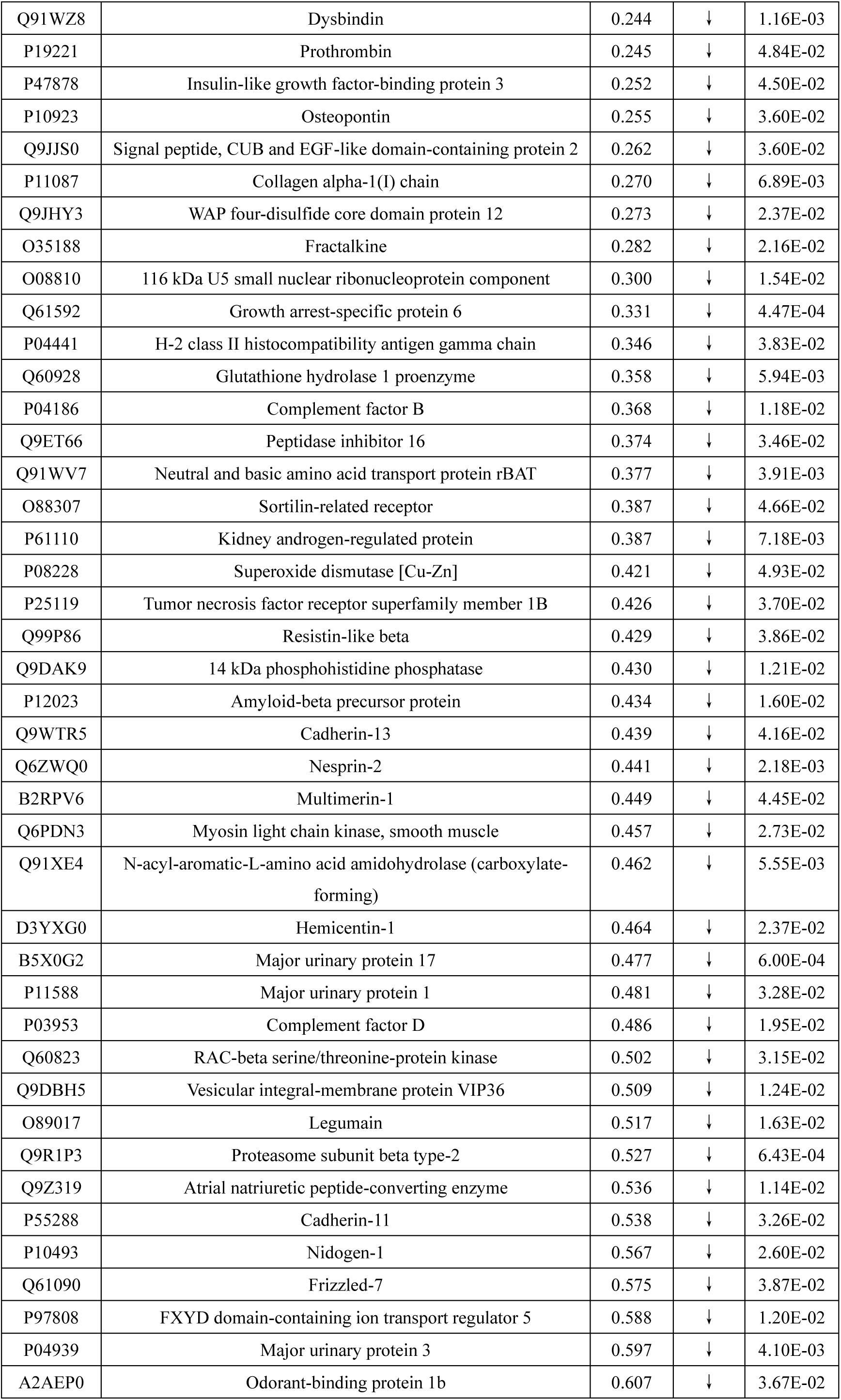
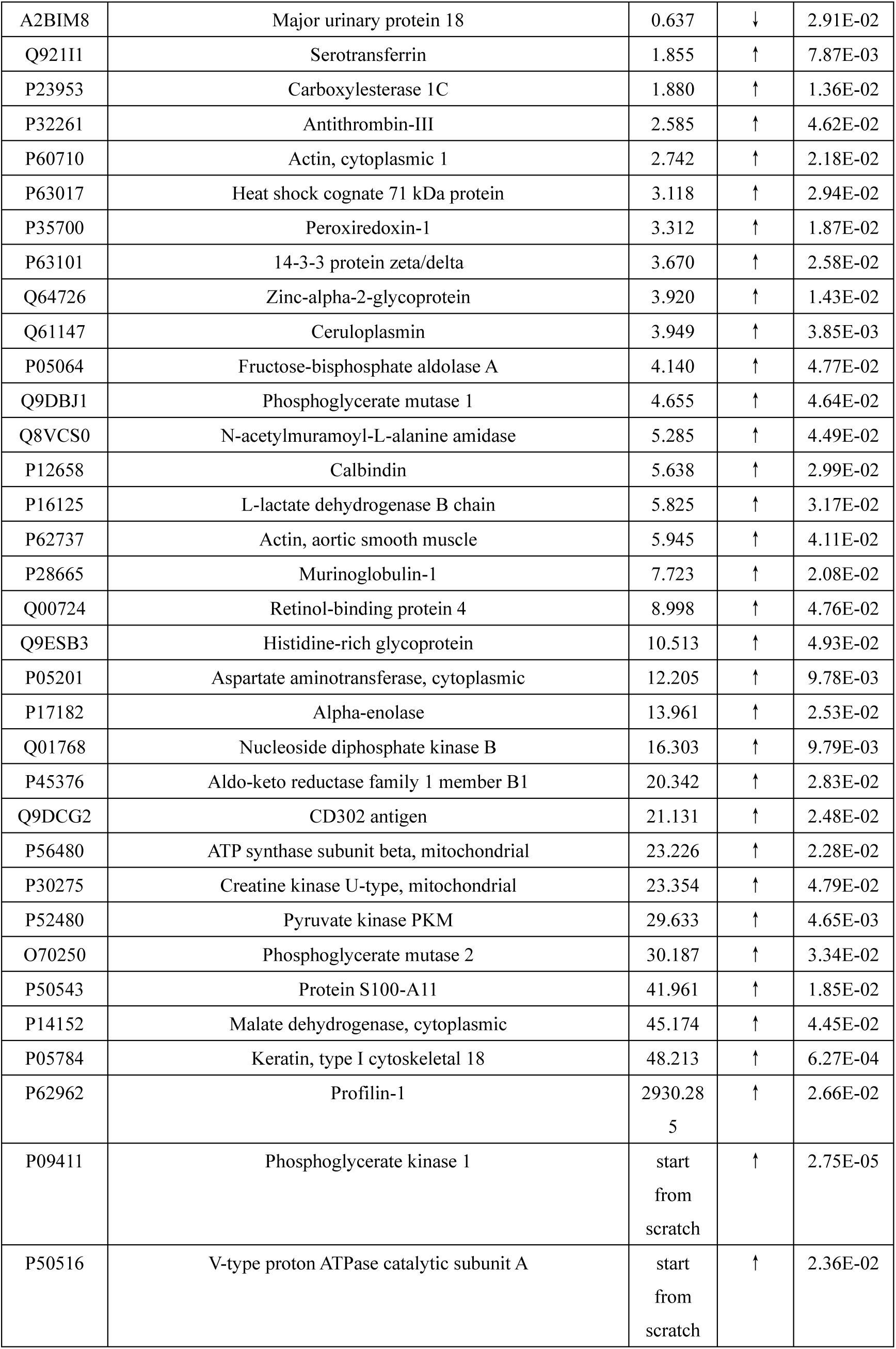
Differential proteins in urine proteome of mice in sucralose experimental group and sucralose control group protein.

#### 3.7.2 Analysis of differential protein function

The 83 identified differential proteins were analysed by DAVID database for molecular functions and biological processes, and the results are shown in Figure 8.

**Figure 8.**
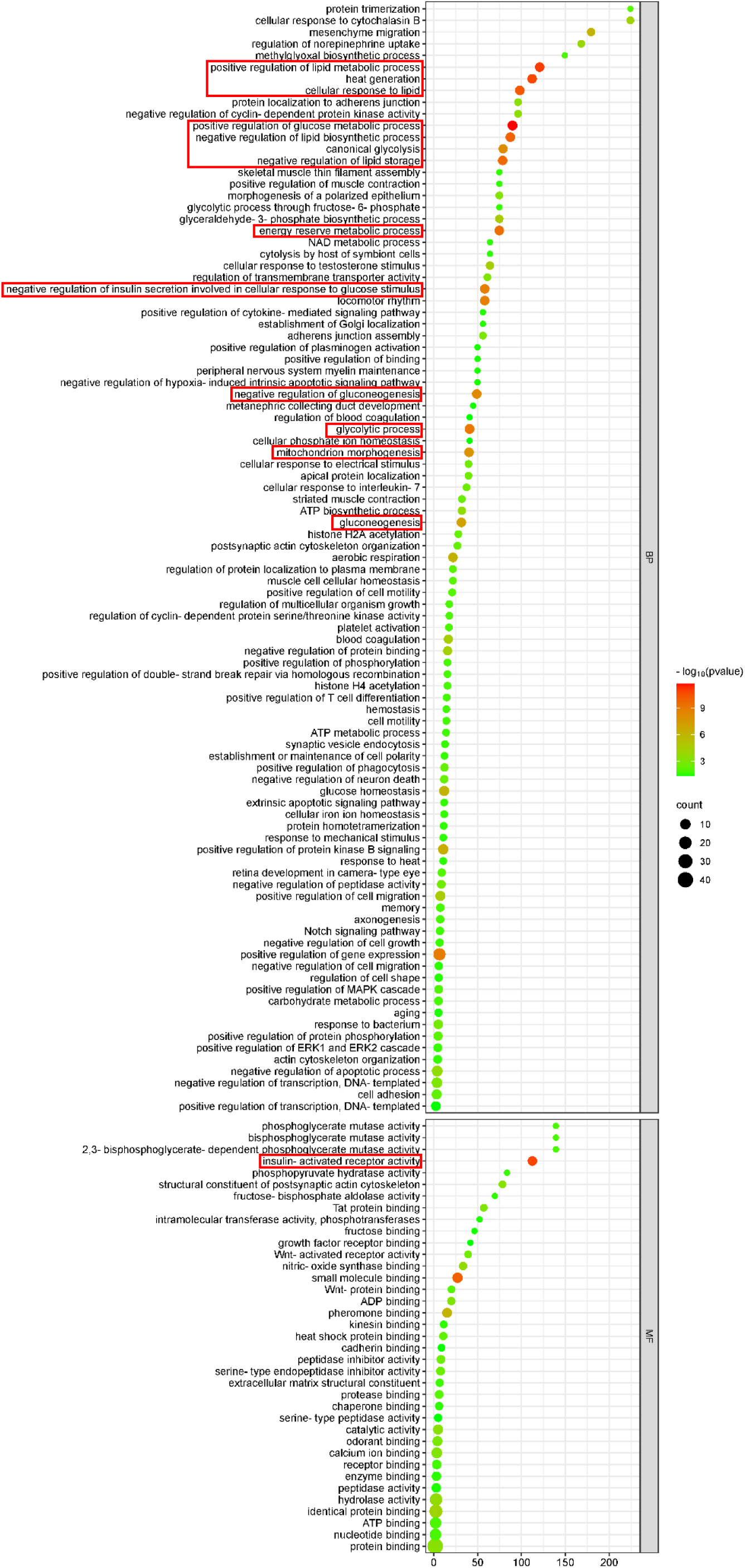
Analysis of molecular functions and biological processes of differential proteins in mice of sucralose group

Sorted by p-value from smallest to largest, among the 28 biological processes ranked in the top 30%, 16 biological processes are involved in a large number of sugar and lipid metabolism and energy production, such as: positive regulation of glucose metabolism, positive regulation of lipid metabolism, heat production, cellular response to lipids, negative regulation of lipid biosynthesis, negative regulation of lipid storage, energy reserve metabolism process, glycolysis process, negative regulation of insulin secretion involved in cellular response to glucose stimulation, etc., which were highly similar to the sucrose group; the molecular functions were also similar to those of the sucrose group, ranked by p-value from the smallest to the largest, in which the first ranked molecular function was insulin receptor activation. Notably, the differential proteins were similarly enriched for neural-related biological processes such as negative regulation of neuronal death, axonogenesis, postsynaptic actin cytoskeleton organisation, synaptic vesicle endocytosis, and maintenance of myelin in the peripheral nervous system, more so than sucrose, stevia glycosides, and anserine, which may be relevant to the preference response of sweet taste perception.

#### 3.7.3 Differential proteins and brain reward circuits

To further investigate whether the urinary proteome differential proteins have brain reward circuit-associated proteins, the functions and biological processes involved in the sucralose group differential proteins were searched by Uniprot to find whether they are related to the key proteins that have been reported to be involved in the brain reward circuits, and the results are shown in Table 11.

**Table 11.**
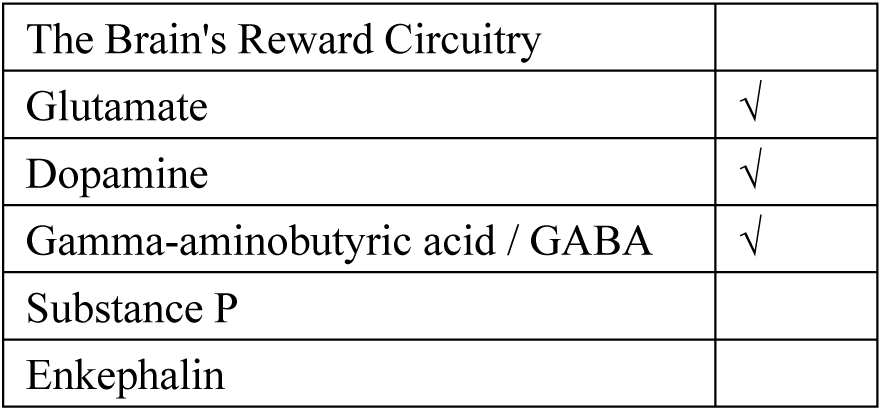
Urinary proteomic differential proteins and brain reward circuits in sucralose group mice.

Mouse urine proteome difference proteins were retrieved with proteins associated with brain reward circuits.

Na(+)/H(+) exchange regulatory cofactor NHE-RF1 functions in dopamine receptor binding and γ-aminobutyric acid transmembrane transport, and is involved in biological processes such as adenylate cyclase-activated dopamine receptor signalling pathway and phospholipase c-activated dopamine receptor signalling pathway.

Glypican-4 is located in regions expressed at glutamatergic synapses, synapses and pre-synaptic membranes, and can participate in the development of the central nervous system, and is involved in biological processes such as the regulation of postsynaptic specialised membranes by neurotransmitter receptor localisation, regulation of presynaptic assembly, and synaptic membrane adhesion.

Dysbindin is located in asymmetric axons, axon cytoplasm, dendritic spines, neuronal cytosol, postsynaptic membrane and other regions of expression, with synaptic vesicle transport, neurotransmitter release, neuronal synaptic growth and affect the glutamatergic release to promote neuronal transmission and viability of the function, involved in the paracentric axon transport, paracentric synaptic vesicle transport, dendritic morphogenesis, glutamatergic neurotransmitter secretion, dopamine receptor signalling pathway regulation, dopamine secretion regulation, synaptic vesicle cytosolic regulation and other biological processes.

Fractalkine is located in the neuronal cytosol and other regions of expression and is involved in biological processes such as inhibitory postsynaptic potentials, negative regulation of glutamate receptor signalling pathways negative regulation of apoptotic processes in hippocampal neurons, neuronal cellular homeostasis, and positive regulation of neuroblast proliferation.

Glutathione hydrolase 1 proenzyme has glutathione hydrolase activity and is involved in glutathione catabolism, glutamate metabolic processes and response to alcohol.Amyloid-beta precursor protein acts as a cell-surface receptor and performs physiological functions on the surface of neurons related to neuronal synapse growth, neuronal adhesion and axonogenesis; it is involved in biological processes such as learning and memory signalling pathways, associative learning and neuronal differentiation. genesis; it is involved in biological processes such as learning and memory, glutamate receptor signalling pathways, associative learning, and neuronal differentiation.

Amino acid transporter heavy chain SLC3A1 is involved in the transmembrane transport of glutamate.

Amyloid-beta precursor protein acts as a cell surface receptor on the surface of neurons to perform physiological functions related to neurite growth, neurite adhesion, and axonogenesis; it is involved in biological processes such as learning and memory, glutamate receptor signalling pathways, associative learning, and neuronal differentiation.

Cadherin-13 is expressed in gamma-aminobutyric acidergic synapses, neural projections and other regions, and is involved in biological processes such as positive regulation of calcium-mediated signalling pathways

Cadherin-11 is expressed in regions such as glutamatergic synapses and is involved in biological processes such as corticospinal tract morphogenesis and regulation of chemical synaptic transmission.

Heat shock cognate 71 kDa protein is expressed in regions of asymmetric axons, axons, dendrites and dendritic spines, glutamatergic synapses, myelin sheaths, postsynaptic membranes, and synaptic vesicles, and is involved in the regulation of postsynaptic organisation, response to odour, slow axonal transport, and other biological processes.

14-3-3 protein zeta/delta is expressed in regions such as glutamatergic synapses and is involved in biological processes such as the regulation of synaptic maturation and recognition of synaptic targets.

Calbindin is expressed in axons, dendrites, dendritic spines, gamma-aminobutyric acidergic synapses, glutamatergic synapses, neuronal cytosol, and other regions, and is involved in biological processes such as the regulation of synaptic plasticity, the regulation of pre and post-synaptic cytoplasmic calcium ion concentrations through calcium binding, the regulation of pre-synaptic cytoplasmic calcium ion concentration, long term memory, short term memory, and so on.

Aspartate aminotransferase, cytoplasmic catalyses the synthesis of glutamate: a major excitatory neurotransmitter of the vertebrate central nervous system and an important regulator of glutamate levels. Acts as a glutamate scavenger in brain neuroprotection. Involved in biological processes such as glutamate synthesis and catabolism.

Profilin-1 is expressed at glutamatergic synapses, pre- and postsynaptic cells, and other regions, and is involved in the regulation of chemical synaptic transmission, synapse maturation, and other biological processes.

### 3.8 Individual analyses of the proteome of the anterior urine of individuals in the sucralose group after sucralose consumption

The urine proteome of five mice after sucralose consumption was compared with the urine proteome before sucralose consumption, respectively, and the condition of screening for differential proteins was FC ≥1.5 or ≤0.67, and the t-test was P<0.05, and the differential proteins shared by five mice were counted, and the results are shown in Figure 9.

**Figure 9.**
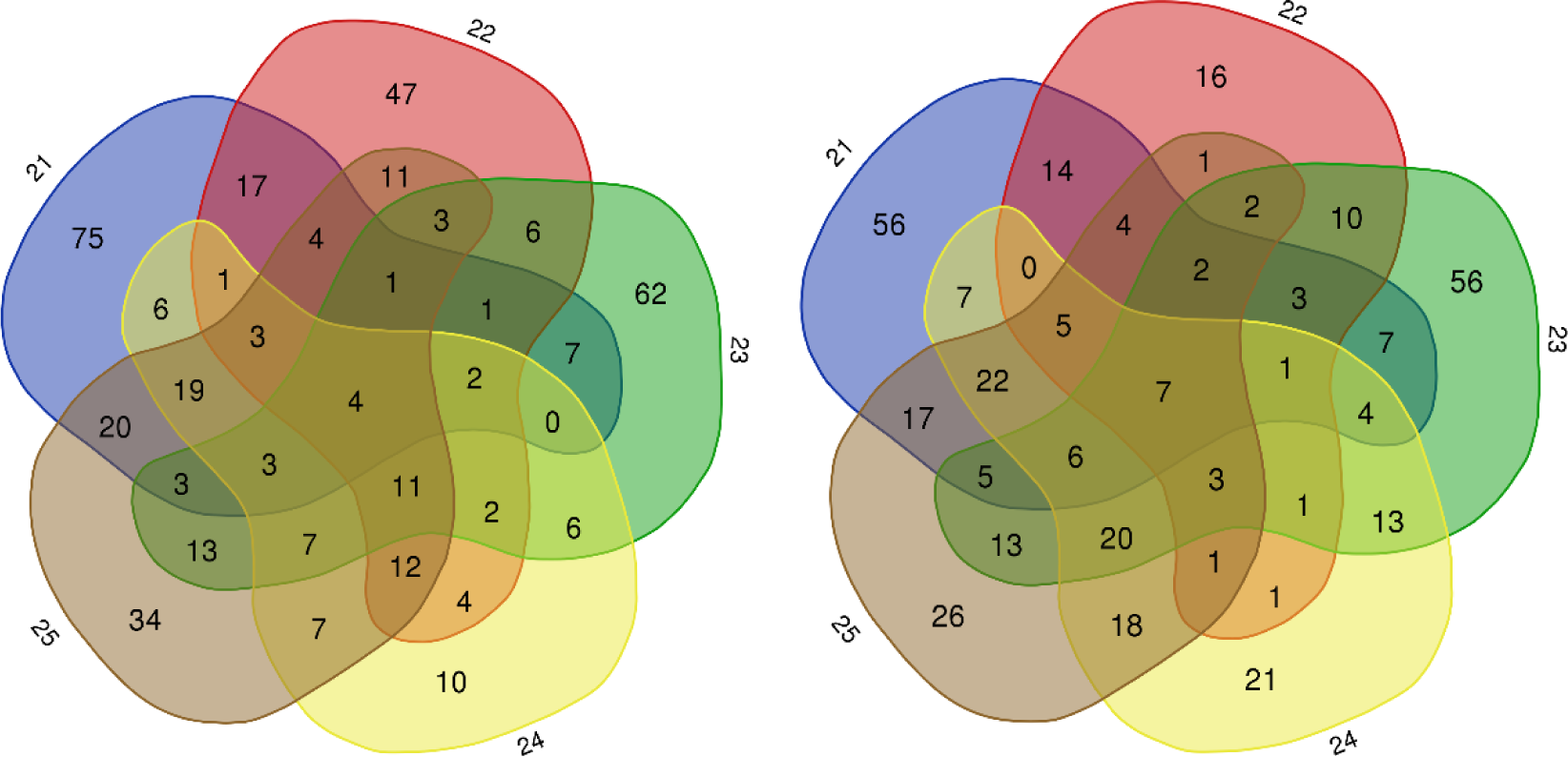
Total up-regulated differential proteins (left) down-regulated differential proteins (right) in front of individuals in the sucralose group after sucralose consumption

There were a total of 4 up-regulated differential proteins and 12 down-regulated differential proteins in 5 mice, as shown in Table 12.

**Table 12.** Pre-shared differential proteins after sucralose consumption by individuals in the sucralose group.

| Uniprot ID | Protein name | Trend |
| --- | --- | --- |
| P05784 | Keratin, type I cytoskeletal 18 | ↓ |
| P10923 | Osteopontin (2AR) | ↓ |
| P41272 | CD27 antigen | ↓ |
| P15626 | Glutathione S-transferase Mu 2 | ↓ |
| P04939 | Major urinary protein 3 | ↓ |
| P70441 | Na(+)/H(+) exchange regulatory cofactor NHE-RF1 | ↓ |
| Q60928 | Glutathione hydrolase 1 proenzyme | ↓ |
| P11087 | Collagen alpha-1(I) chain | ↑ |
| P09411 | Phosphoglycerate kinase 1 | ↑ |
| P12658 | Calbindin | ↑ |
| Q02596 | Glycosylation-dependent cell adhesion molecule 1 | ↑ |

The functions of the proteins and the biological processes involved were retrieved by Uniprot, in which Na(+)/H(+) exchange regulatory cofactor NHE-RF1 has the functions of dopamine receptor binding and γ-aminobutyric acid transmembrane transport, and is involved in the adenylyl cyclase-activated dopamine receptor signalling pathway, phospholipase c-activated dopamine receptor signalling pathway Calbindin is expressed in axons, dendrites, dendritic spines, gamma-aminobutyric acidergic synapses, glutamatergic synapses, neuronal cytosol, etc. It is involved in the regulation of synaptic plasticity, the regulation of pre and post-synaptic cytoplasmic calcium concentration through calcium binding, the regulation of pre-synaptic cytoplasmic calcium concentration, long term memory, short term memory and other biological processes. and other biological processes. This may be related to the sweet taste preference response and brain reward circuits.Glutathione hydrolase 1 proenzyme has glutathione hydrolase activity and is involved in the catabolism of glutathione, the metabolic process of glutamate and the response to alcohol.

The urine proteomic differential proteins of individual mice before and after sucralose consumption were also searched for brain reward circuit-associated proteins that were present in the individual mouse urine proteomic differential proteins, as shown in Table 13.

**Table 13.**
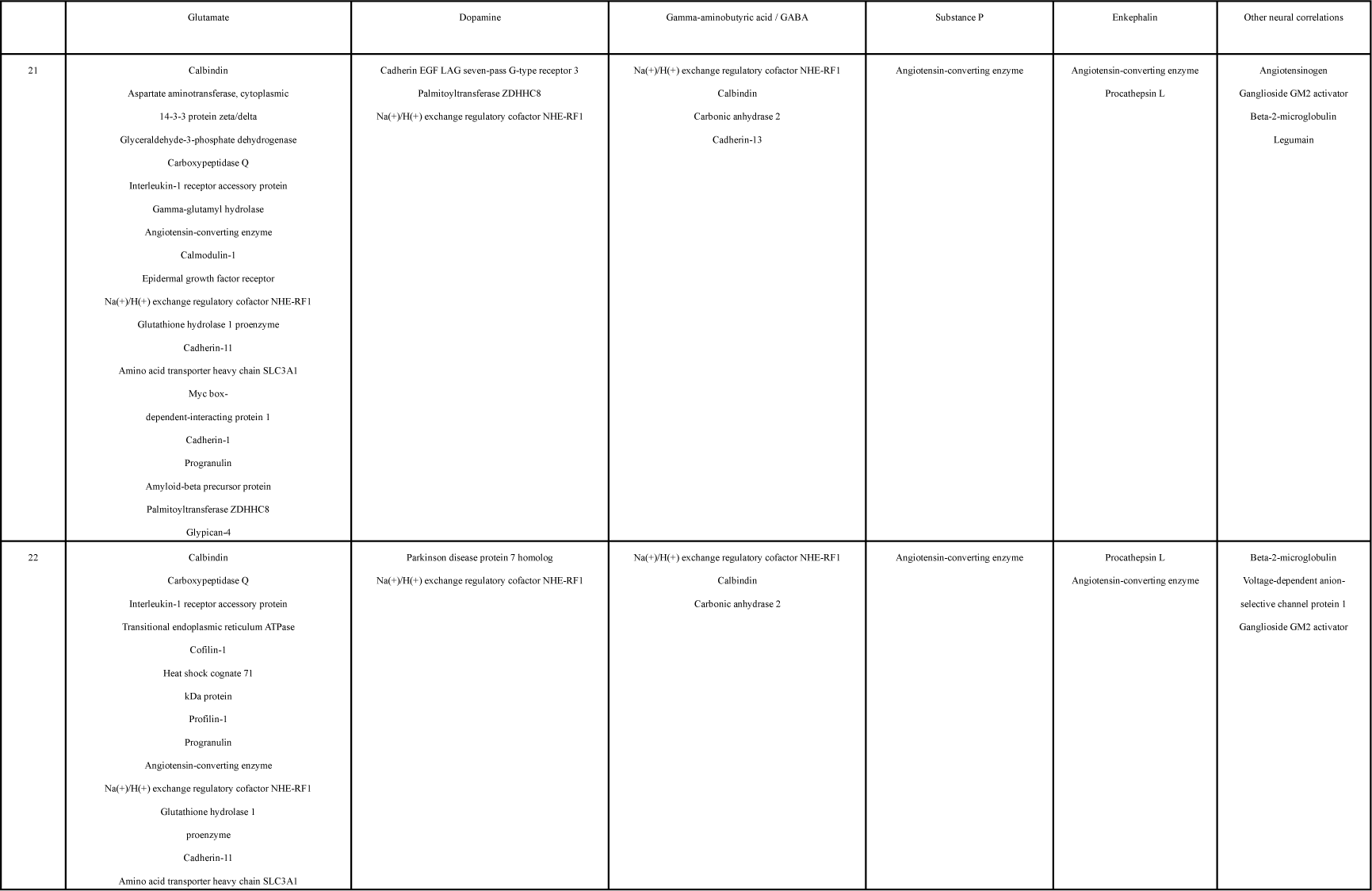
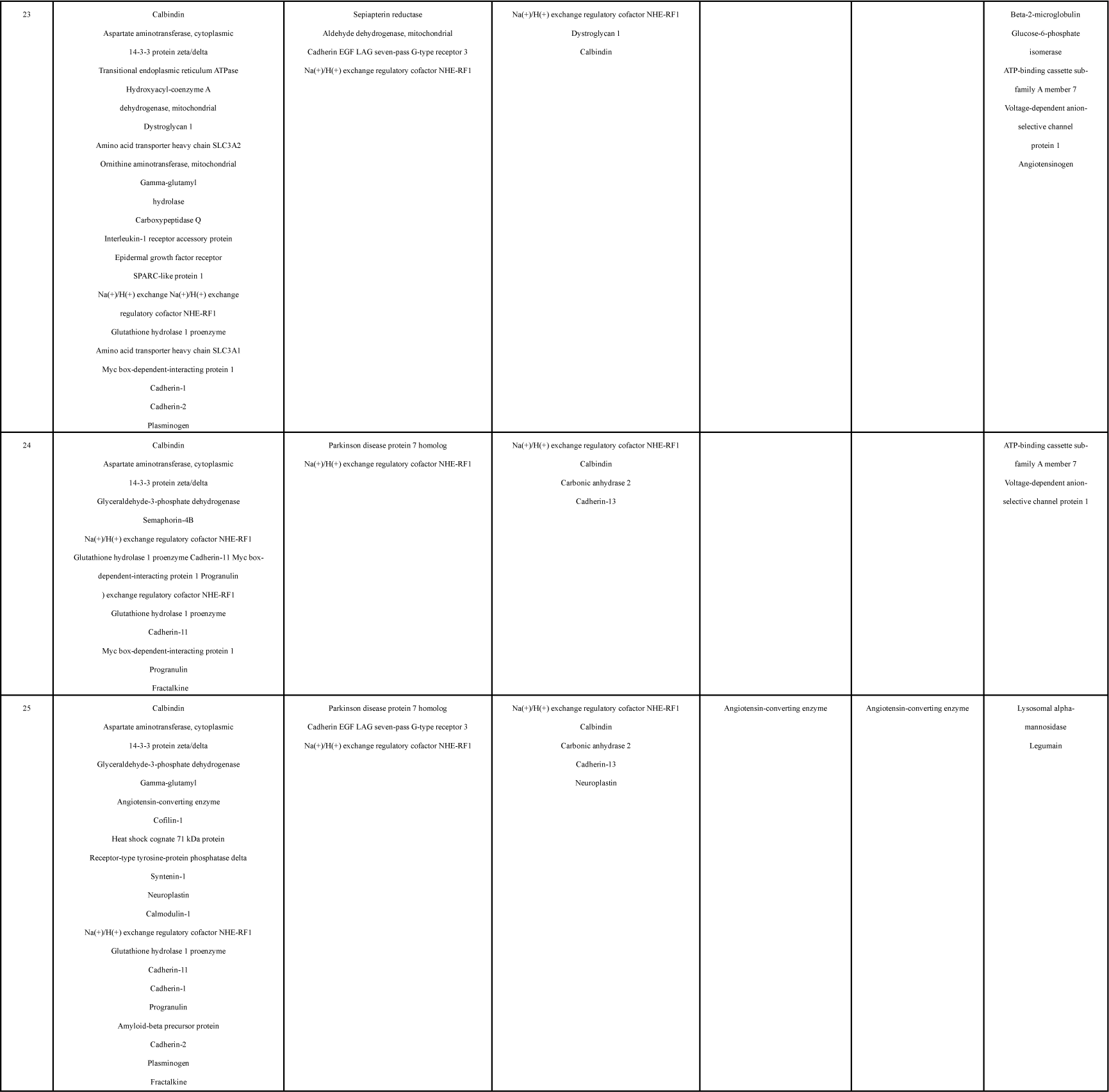
Proteins associated with brain reward circuits in the proteome difference proteins of the anterior urine proteins of individuals in the sucralose group after sucralose consumption.

## 4 Cross-sectional comparative analysis of sweetening substances

### 4.1 Comparison of differential proteins analysed in groups

The common differential proteins for the four sweet substances in the group analysis are shown in Figure 10.

**Fig. 10.**
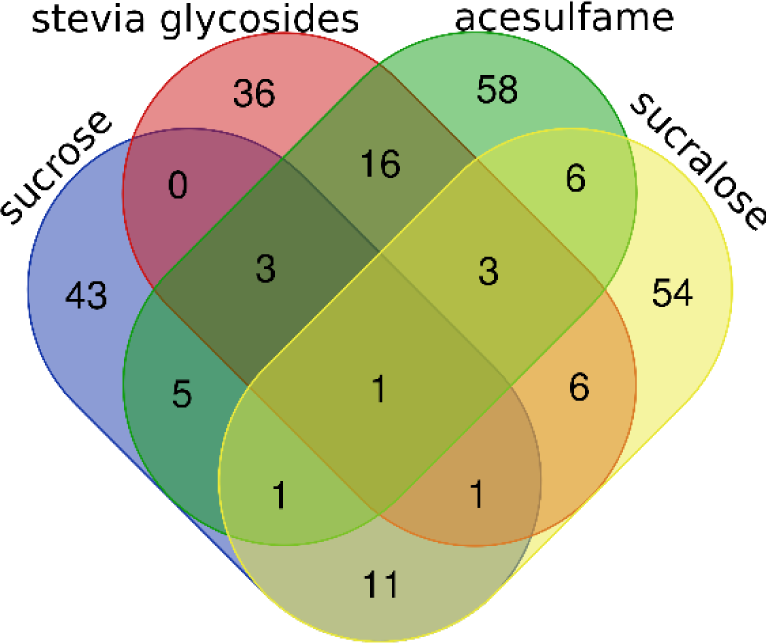
Shared differential proteins analysed in groups of four sweet taste substances

Compared to the sucrose group, the steviol glycoside group shared 5 differential proteins with the sucrose group, the acesulfame group shared 10 differential proteins with the sucrose group, and the sucralose group shared 15 differential proteins with the sucrose group. Compared to stevia glycosides, the acesulfame group shared 23 differential proteins with the stevia glycosides group and the sucralose group shared 11 differential proteins with the stevia glycosides group. Compared with acesulfame, the sucralose group had a total of 11 differential proteins with the acesulfame group. The Acesulfame group had the most total differential proteins when analysed as a group with the Stevia glycosides group, followed by the Sucralose group with the Sucrose group, and the Stevia glycosides group had the least total differential proteins when analysed as a group with the Sucrose group. The specific proteins are shown in Exhibit 1.

The biological processes common to the four sweeteners in the group analysis are shown in Figure 11.

**Fig. 11.**
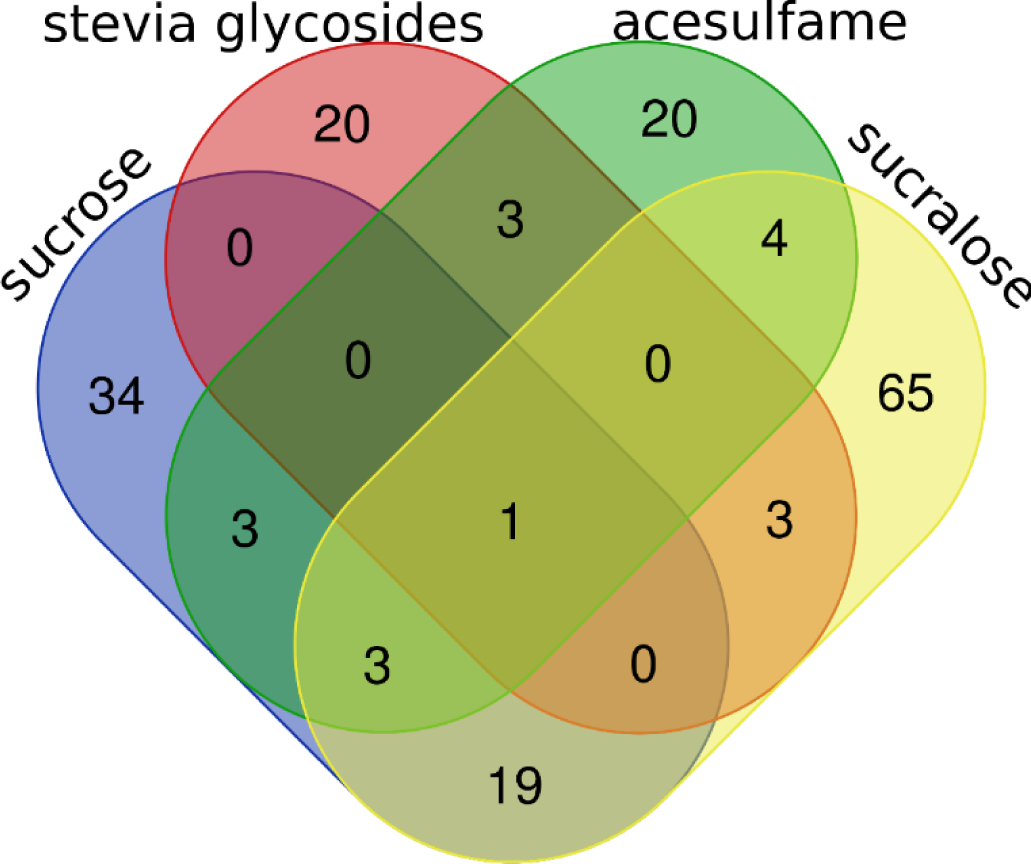
Shared biological processes in the group analysis of the four sweet taste substances

Compared to the sucrose group, the steviol glycoside group shared 1 biological process with the sucrose group, the acesulfame group shared 7 biological processes with the sucrose group, and the sucralose group shared 23 biological processes with the sucrose group. Compared to stevia glycosides, the acesulfame group shared 4 biological processes with the stevia glycosides group and the sucralose group shared 4 biological processes with the stevia glycosides group. Compared to acesulfame, the sucralose group shared 8 biological processes with the acesulfame group. The sucralose group shared the most biological processes and was much higher than the other groups when analysed as a group with sucralose, while the steviol glycoside group shared the fewest differential proteins when analysed as a group with sucralose. It is worth noting that the steviol glycoside group also had fewer shared biological processes with other sweet substances, which may indicate that steviol glycosides affect the organism differently from other sweeteners. The specific biological processes are shown in Exhibit 2.

The biological processes of differential protein enrichment in the four sweet substance groups were all related to the nervous system, although the specific biological processes involved were different and the changes were not the most significant among all the biological processes, but this was likely to be related to the preference response of sweetness sensation; the biological processes of differential protein enrichment in the sucrose group, the acesulfame group and the sucralose group were mainly involved in the glucose and lipid metabolism, the production of energy, etc., with few changes related to the nucleosome The biological processes of differential protein enrichment in the sucrose and sucralose groups mainly involved glycolipid metabolism and energy production, with few changes related to nucleosome assembly and gene expression, whereas the biological processes of differential protein enrichment in the stevia glycosides group mainly involved nucleosome assembly, gene expression and cell division, with few changes related to energy metabolism, which is likely to indicate that the organismal responses induced by the consumption of stevia glycosides are different from those of the other three sweet taste substances.

The molecular functions of the shared differential proteins of the four sweeteners in the group analysis are shown in Figure 12.

**Fig. 12.**
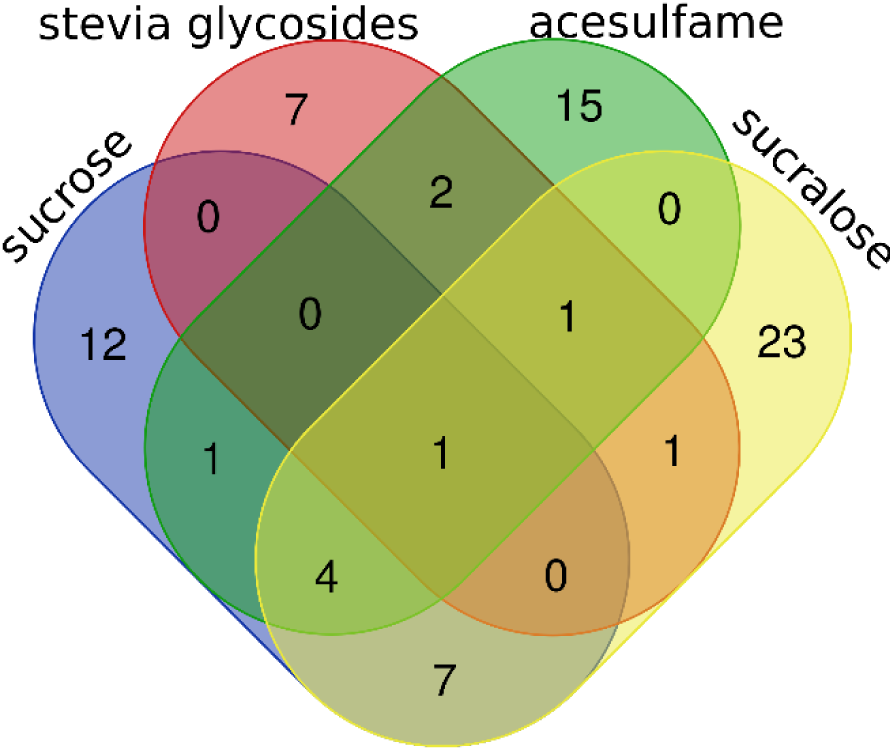
Molecular functions of shared differential proteins analysed in groups of four sweet taste substances

Compared to the sucrose group, the steviol glycoside group shared 1 shared differential protein molecular function with the sucrose group, the acesulfame group shared 4 shared differential protein molecular functions with the sucrose group, and the sucralose group shared 12 shared differential protein molecular functions with the sucrose group. Compared with stevia glycosides, the acesulfame group shared 4 shared differential protein molecular functions with the stevia glycosides group, and the sucralose group shared 3 shared differential protein molecular functions with the stevia glycosides group. Compared with acesulfame, the sucralose and acesulfame groups shared six common differential protein molecular functions. The sucralose and sucralose groups had the highest number of shared differential protein molecular functions when analysed as a group and were much higher than the other groups, while the stevia glycosides and sucralose groups had the lowest number of shared differential protein molecular functions when analysed as a group. Notably, the steviol glycoside group had the least shared differential protein molecular function with other sweet substances. The specific biological processes are shown in Exhibit 3.

The significant presence of insulin receptor activation, which is directly related to blood glucose changes, in the molecular functions of the differential proteins in the sucrose and sucralose groups, but not in the stevia glycosides or acesulfame groups, may indicate that sucralose is more likely to cause blood glucose fluctuations like sucrose when consumed, relative to stevia glycosides and acesulfame.

All four sweet substance group difference proteins in the group analysis contained proteins associated with brain reward circuit processes, as shown in Table 14.

**Table 14.**
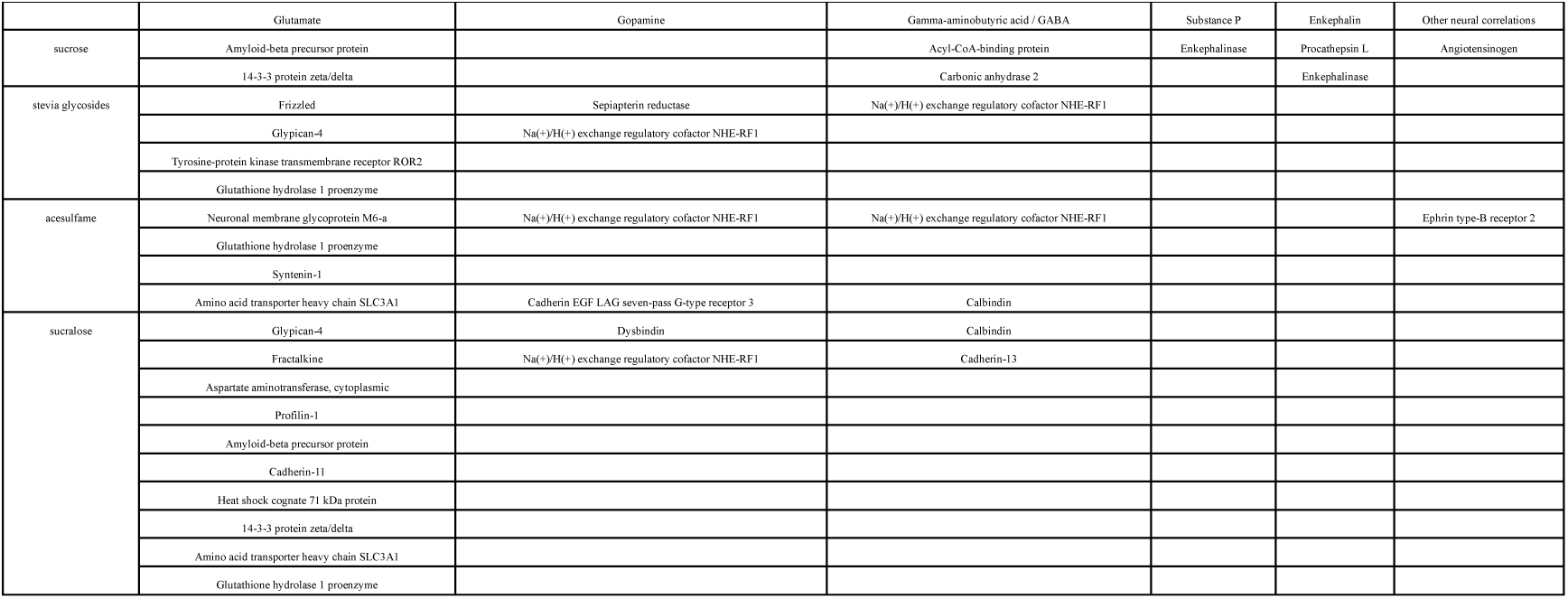
Differential proteins with brain reward circuits in group-wise analyses of four sweet taste substances.

The sucrose and sucralose groups involved more types of brain reward circuit-related proteins than the other groups, which may indicate that the brain reward circuits induced by sucrose and sucralose are more complex and may be more addictive.

None of the four sweet substance group difference proteins appeared to be associated with sweet sensory perception, and it is hypothesised that it may be that proteins for sweet sensory perception do not appear in the urinary proteome, but proteins associated with brain reward circuits appeared in the urinary proteome difference proteins.

Based on the results of the group analysis above, it is tentatively inferred that among the four sweet substances, sucralose and sucrose caused the most similar changes in the organism, and steviol glycosides were the furthest away from sucrose, and the changes caused by steviol glycosides were different from those caused by other sweet substances. 4 sweet substances all caused the appearance of different proteins in the urinary proteome that were related to the brain reward circuits, and only sucrose, acesulfame, and sucralose caused a large number of changes in metabolic processes in the urinary proteome. Sucrose induced a large number of metabolic process changes in the urinary proteome.

### 4.1 Comparison of individual analyses of shared differential proteins

The common differential proteins shared by the four sweeteners in the individual analyses are shown in Figure 13.

**Fig. 13.**
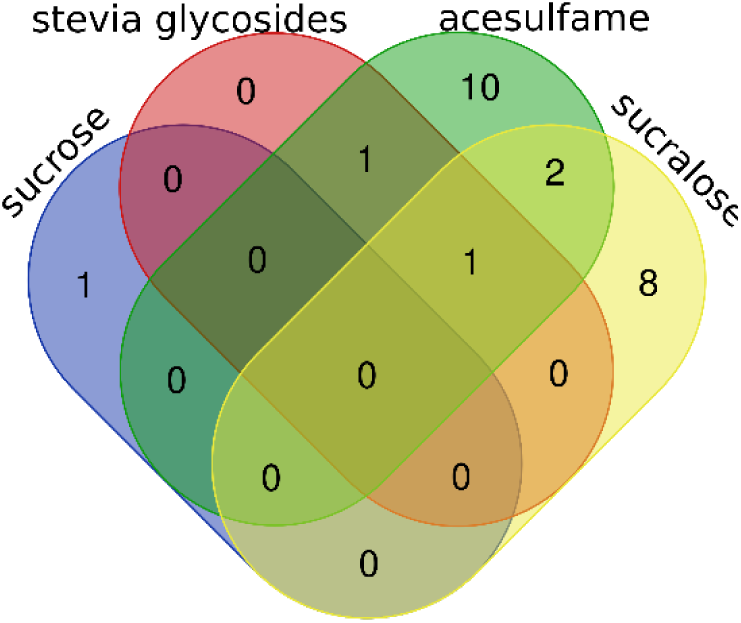
Shared differential proteins common to the individual analyses of the four sweet substances

When analysed individually, the sucrose group only had shared differential proteins related to brain reward circuits, the steviol glycoside group had shared differential proteins related to mRNA metabolism and brain reward circuits, and the shared differential proteins in the acesulfame and sucralose groups were mainly various hydrolytic enzymes involved in metabolic processes, as well as shared differential proteins related to brain reward circuits.

Compared to the sucrose group, the steviol glycoside group had 0 shared differential proteins with the sucrose group, the acesulfame group had 0 shared differential proteins with the sucrose group, and the sucralose group had 0 shared differential proteins with the sucrose group. Compared to stevia glycosides, there were 2 shared differential proteins in the acesulfame group and 1 shared differential protein in the sucralose group and the stevia glycosides group. Compared with acesulfame, the sucralose group had 2 shared differential proteins with the acesulfame group. Among them, the sucralose group and acesulfame group shared the most shared differential proteins when analysed individually, and the 2 shared differential proteins were related to brain reward circuits, and the trends of these two proteins were the same. The specific shared differential proteins are shown in Exhibit 4.

These results may indicate that the presence of proteins related to brain reward circuits in the urinary proteome differential proteins caused by the four sweet substances is relatively stable between individuals. The presence of proteins related to glycolipid metabolism, etc., in the urinary proteome differential proteins caused by sucrose, acesulfame, and sucralose was not relatively stable between individuals in the sucrose group, whereas it was not relatively stable between individuals in the acesulfame and sucralose groups.

## 5 Summary

Group analysis of the differential proteins in the proteome of the anterior urine of mice after autonomous sucrose consumption and the biological processes in which they are enriched, most of them are related to glycolipid metabolism, and a small part of them are related to the biological processes of the nervous system, and the differential proteins include proteins related to the brain reward circuits but not to the perception of sweetness; single analysis of the common differential proteins in the proteome of the anterior urine of mice after autonomous sucrose consumption has proteins related to the brain reward circuit.

The biological processes of differential proteins and their enrichment in the proteome of anterior urine after autonomous stevia glycosides consumption in mice were analysed in groups, most of which were related to nucleosome assembly, gene expression and cell division, and a small part of which were related to the biological processes of the nervous system, and the differential proteins had proteins related to brain reward circuits but not to sweet taste perception; the single analysis of the anterior urine of mice after autonomous stevia glycosides consumption in mice The shared differential proteins of the proteome have proteins related to brain reward circuits.

Unsupervised cluster analysis can distinguish the total urine proteome of mice before and after autonomous consumption of acesulfame, and analyse the differential proteins of the urine proteome of mice before and after autonomous consumption of acesulfame in groups, and their enriched biological processes, which are partly related to glucose and lipid metabolism, but also related to the nervous system, and the differential proteins include proteins related to the brain reward circuits, but not related to the sweetness sensation. The common differential proteins in the anterior urine proteome of single analysed mice after autonomous consumption of acesulfame have proteins related to metabolism and brain reward circuits.

When analysed in groups, most of the differential proteins in the proteome of the anterior urine of mice after autonomous sucralose consumption and the biological processes in which they are enriched are related to glucose and lipid metabolism, and there are also biological processes related to the nervous system, and the differential proteins include proteins related to brain reward circuits but not to sweet taste perception; when analysing the common differential proteins in the proteome of the anterior urine of mice after autonomous sucralose consumption in singles, the proteins have the following characteristics proteins associated with metabolism and brain reward circuits.

The results of the cross-sectional comparisons of the four sweeteners are summarised.Of the four sweeteners, sucralose and sucrose induced the most similar changes in the organism, and steviol glycosides were the furthest removed from the changes induced by sucrose; sucrose, acesulfame, and sucralose induced similar changes in the organism, whereas steviol glycosides induced changes that were different from those of the other sweeteners.The presence of the four sweeteners in the differentially occurring proteins of the urinary proteome Proteins associated with brain reward circuits are relatively consistently present across individuals. The presence of proteins related to glycolipid metabolism in the urinary proteome differential proteins caused by sucrose, acesulfame, and sucralose was not relatively stable between individuals in the sucrose group, whereas the presence of proteins related to glucose and lipid metabolism in the acesulfame and sucralose groups was not relatively stable between individuals in the sucrose and sucralose groups.

This is shown in table 15.

**Table 15.**
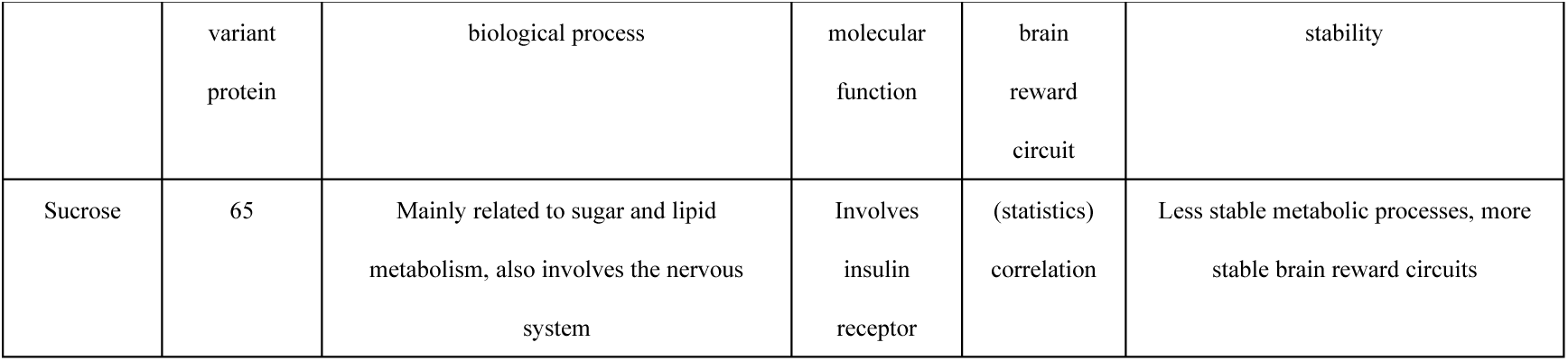
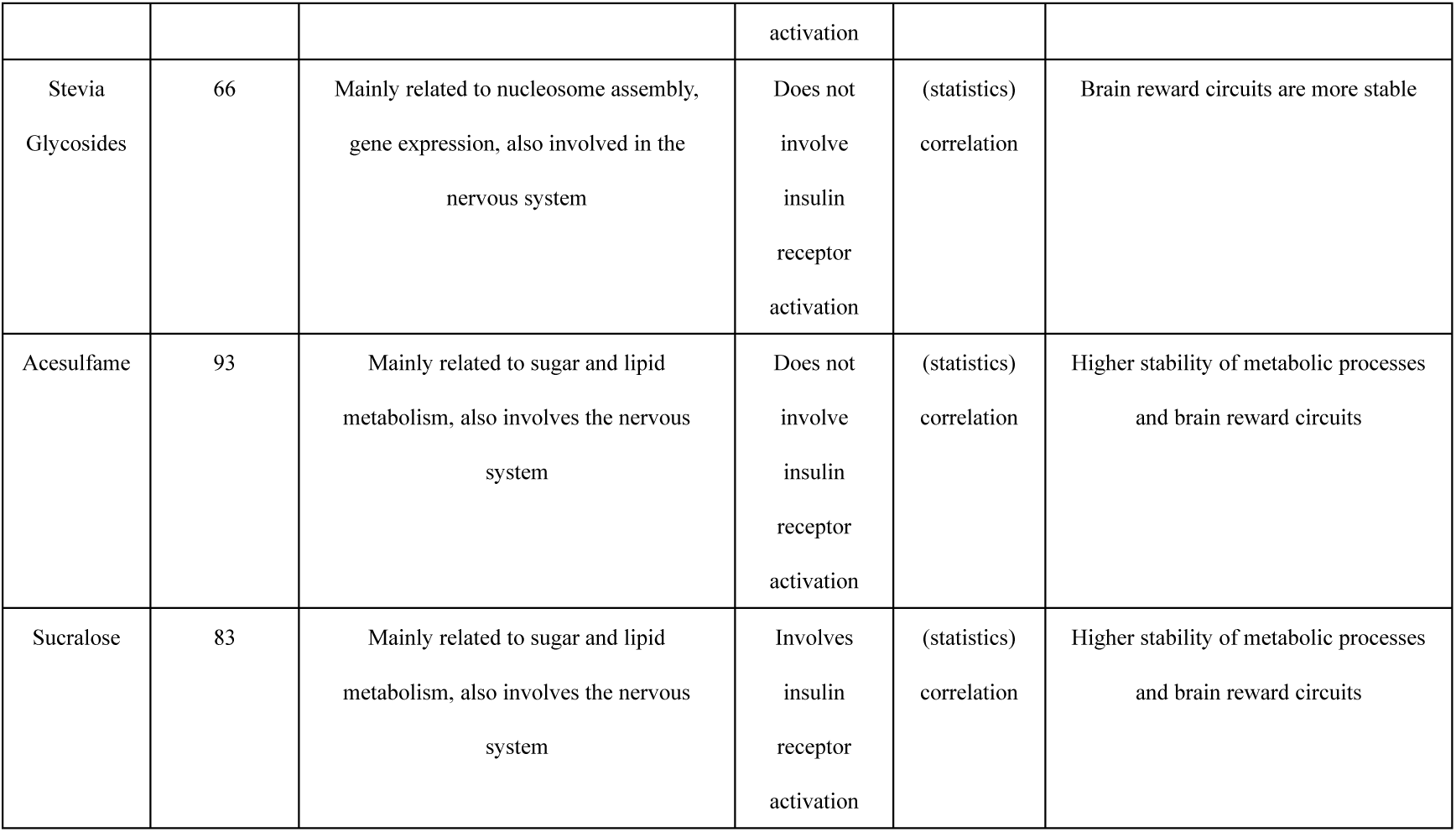
Cross-sectional comparison of urinary proteomic differential proteins of the four sweet substances.

## 6 Discussion

After mice autonomously consumed different sweet substances, the urine proteome changed immediately and in a very rapid response. And the changes caused by different sweet substances are not the same, and the changes are related to metabolism, brain reward circuits and other processes. Previous studies on the effects of different sweeteners on the body and customer preferences for different sweeteners required long-term, large sample sizes for consumption and surveys, which required a large amount of manpower and material resources, whereas studies through urine proteins can greatly shorten the time and cost of the studies. The present study demonstrates the potential of urine proteome in studying the effects of various sweeteners on the body, providing a new approach to urinary proteomics research to explore the development of safer, more stable and more customer-appealing non-nutritive sweeteners. Further experiments may consider using more sweet substances, expanding the sample size of experimental animals or collecting human samples for the study; it also reflects the sensitivity of the urinary proteome and opens up a new field for the exploration of the function of the urinary proteome.

---

**Table 1.**
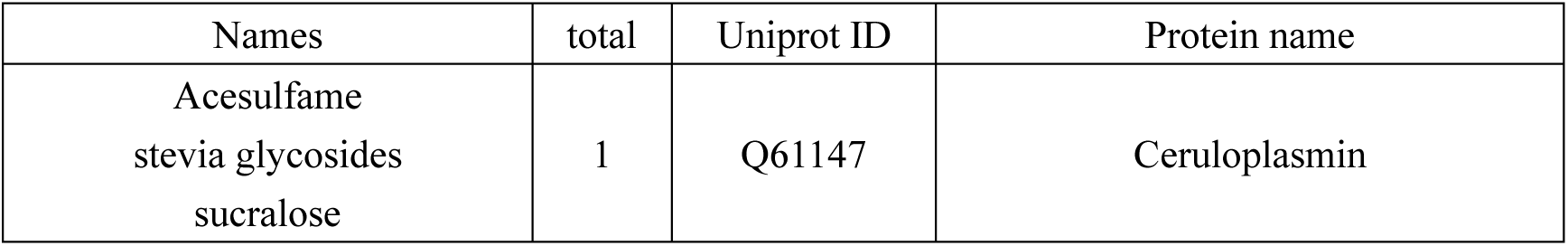
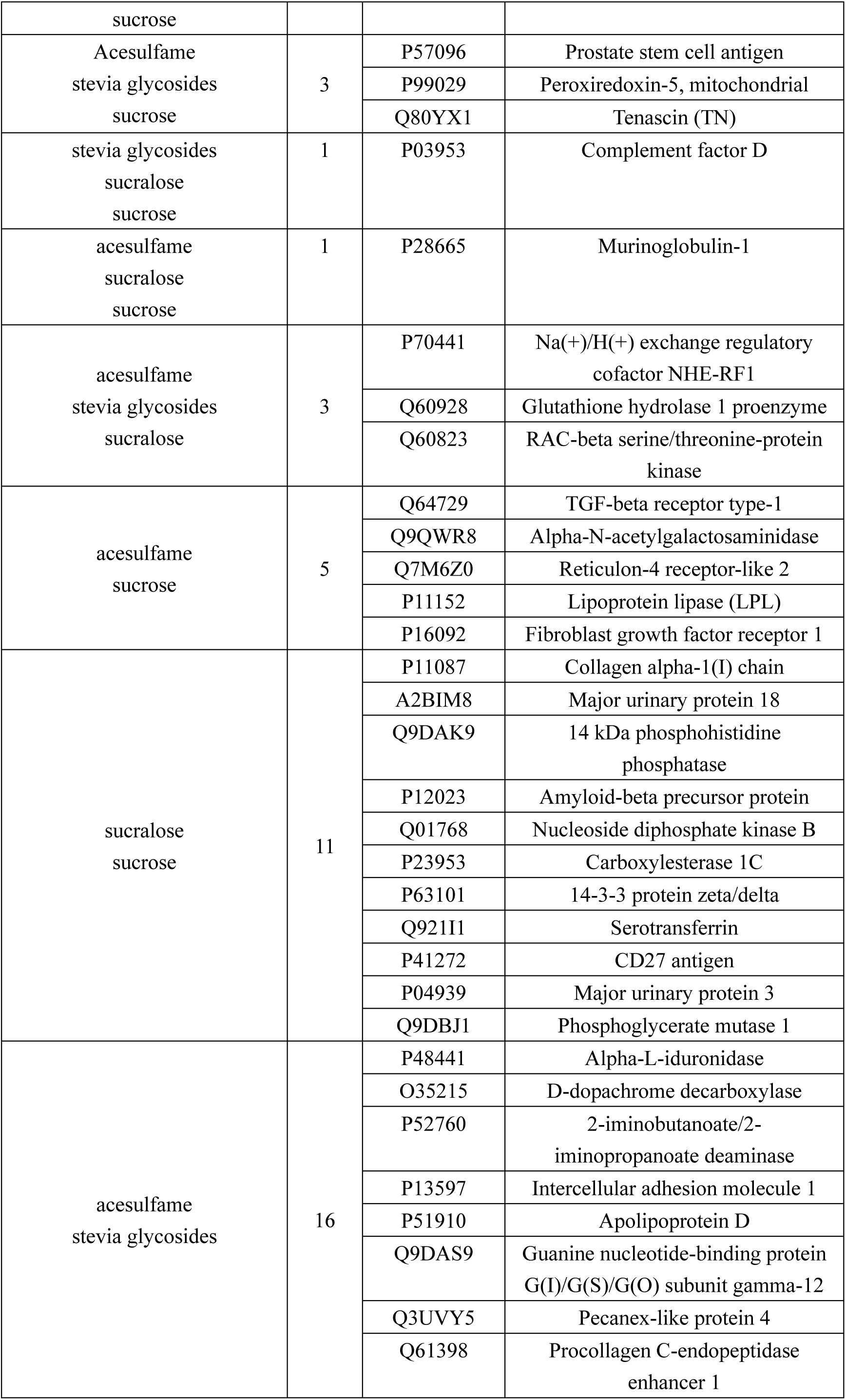
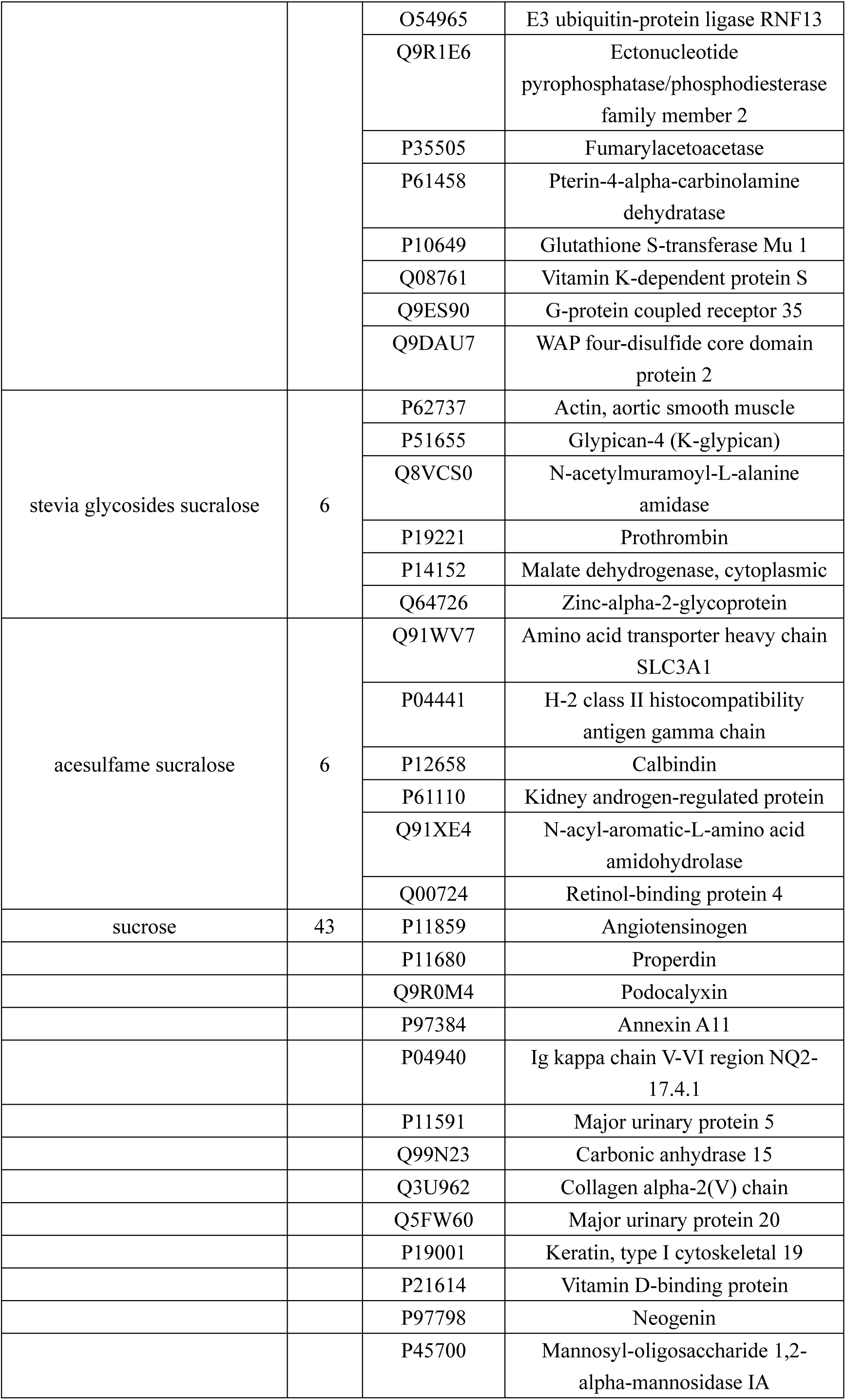
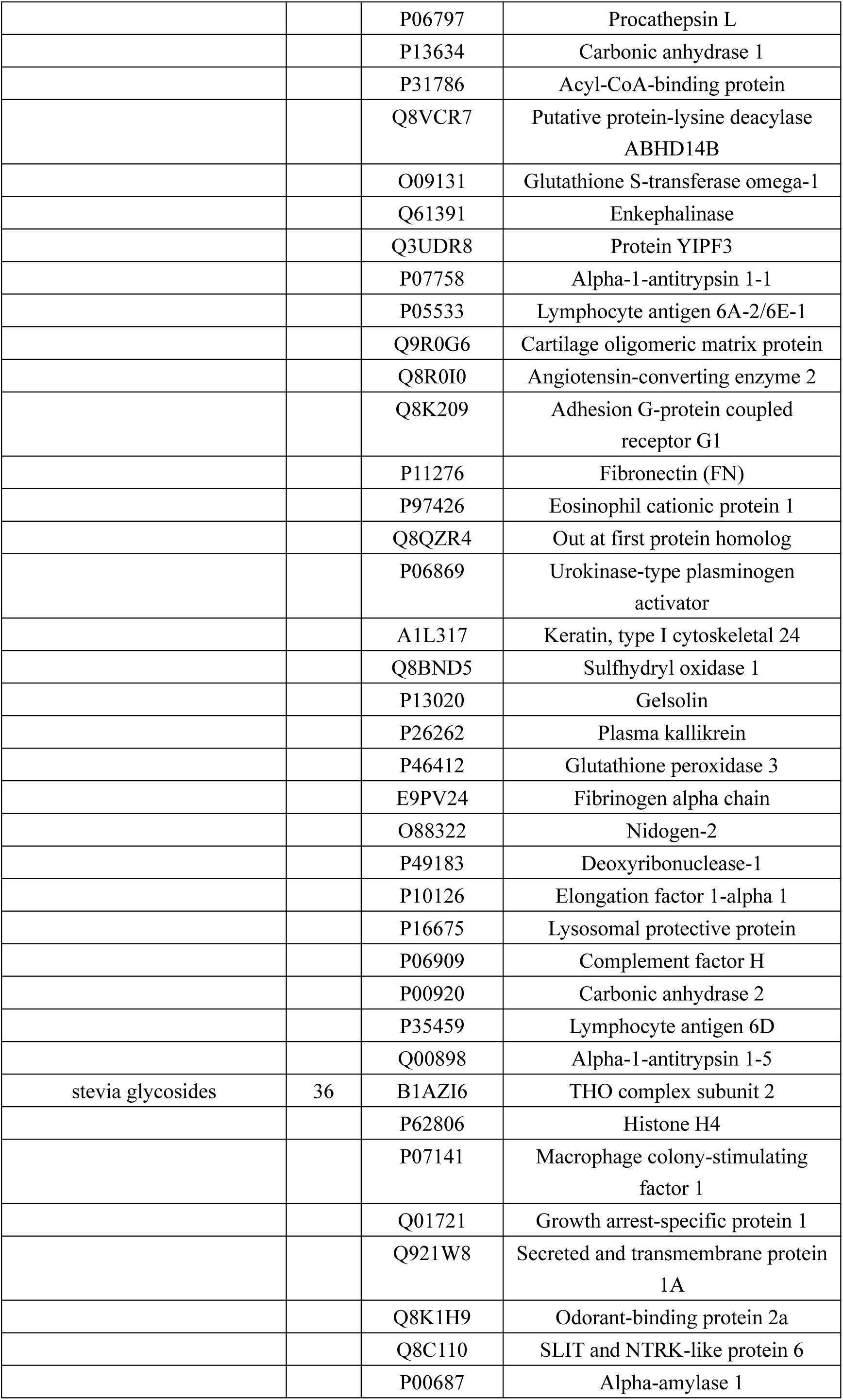
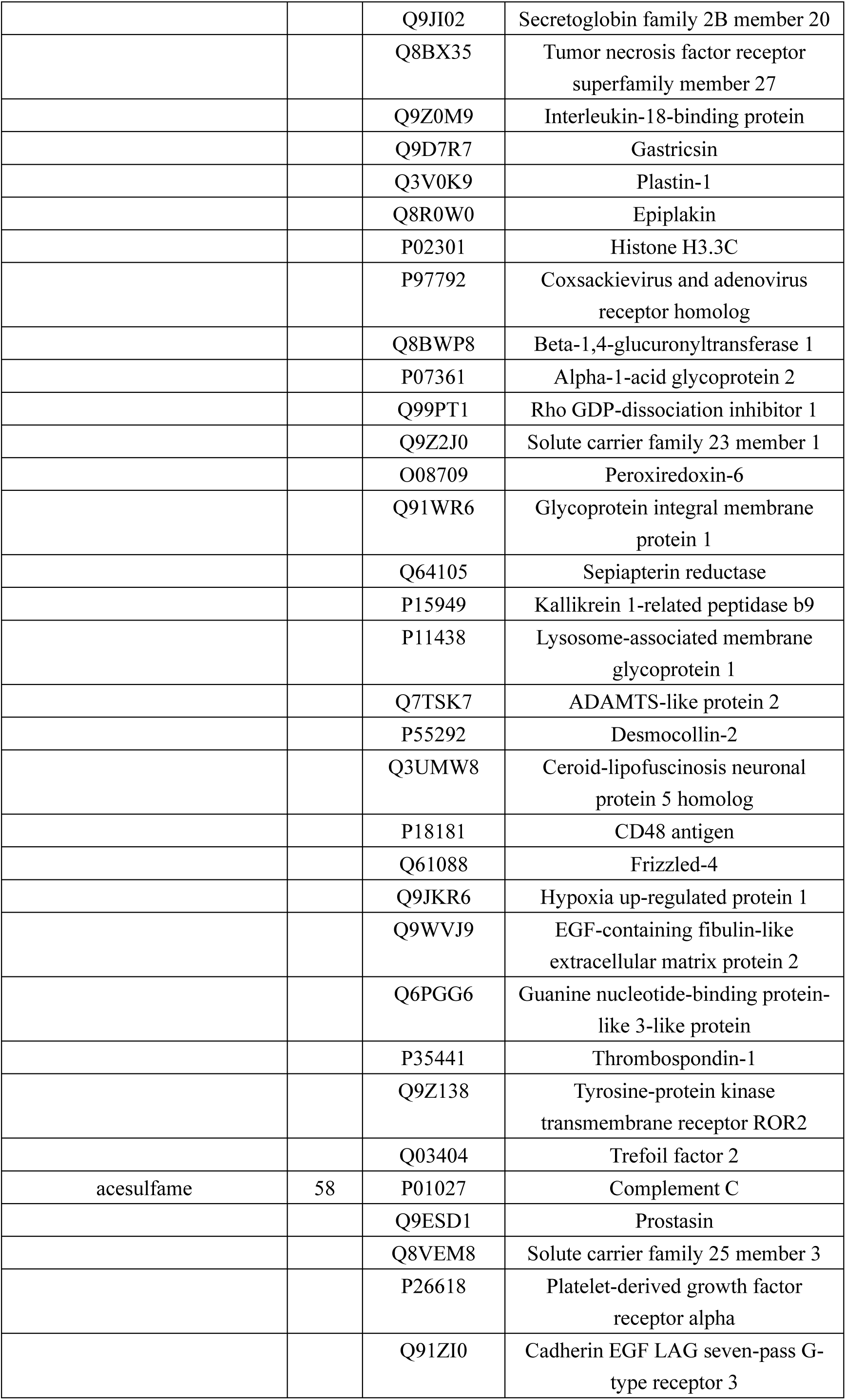
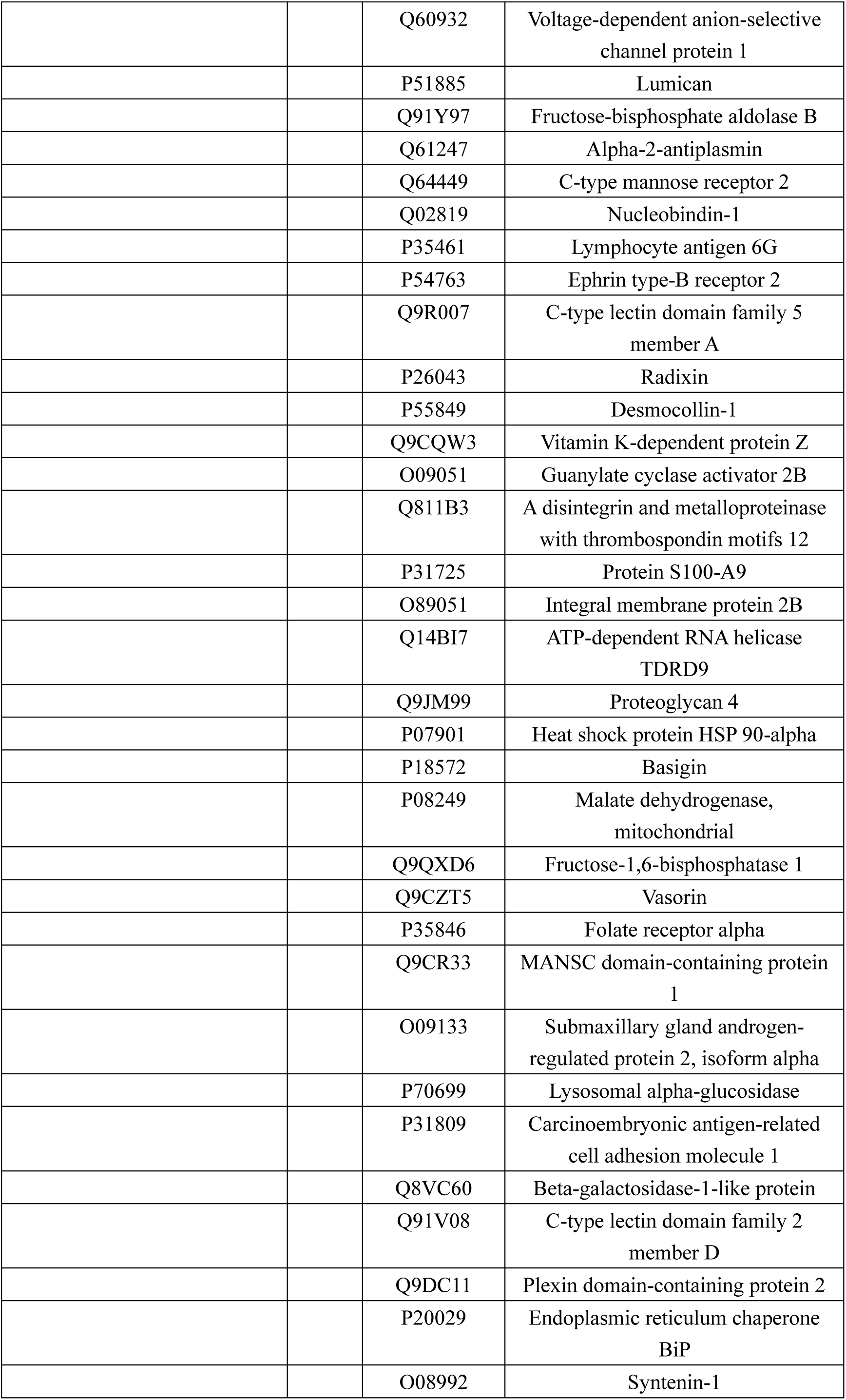
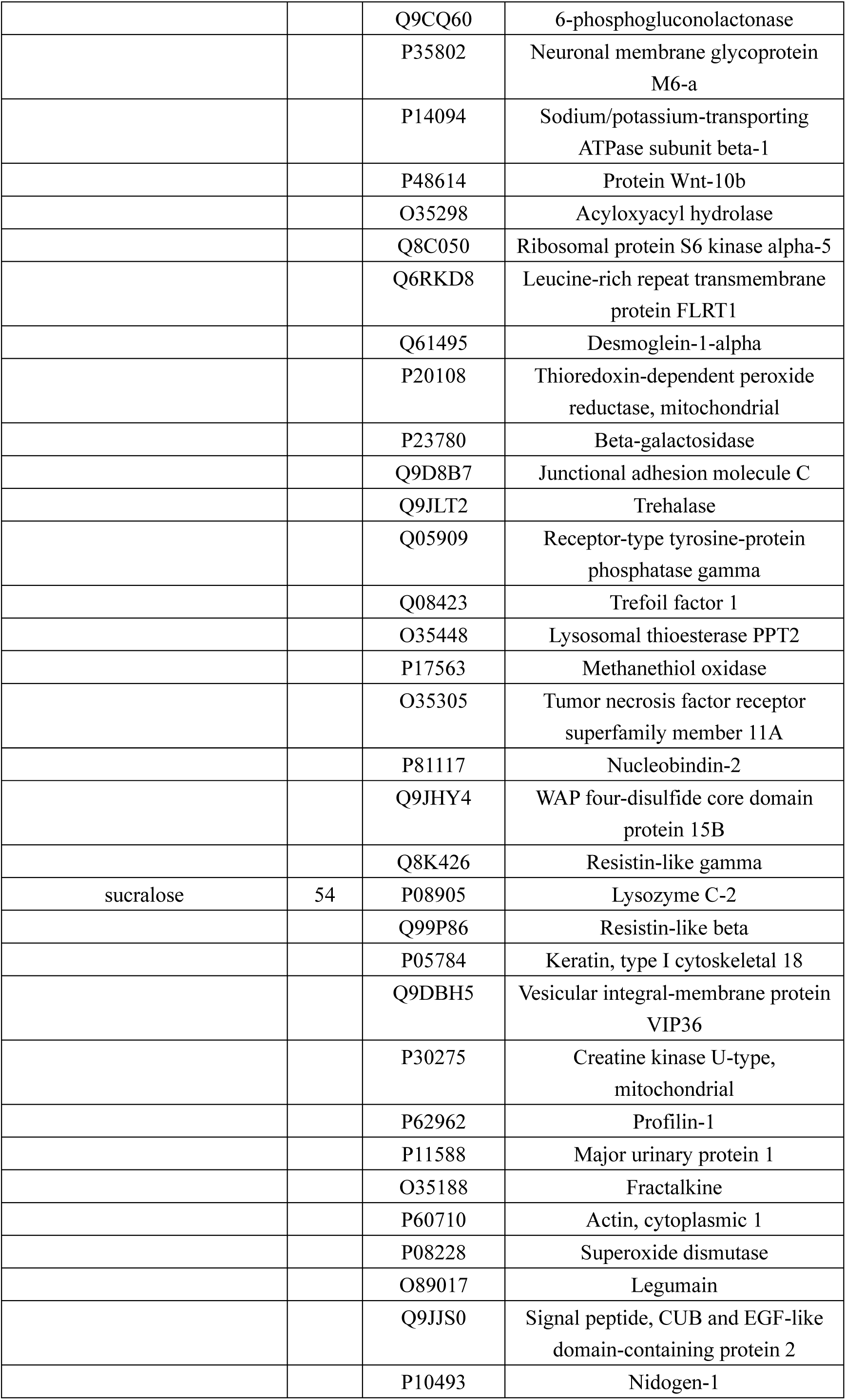
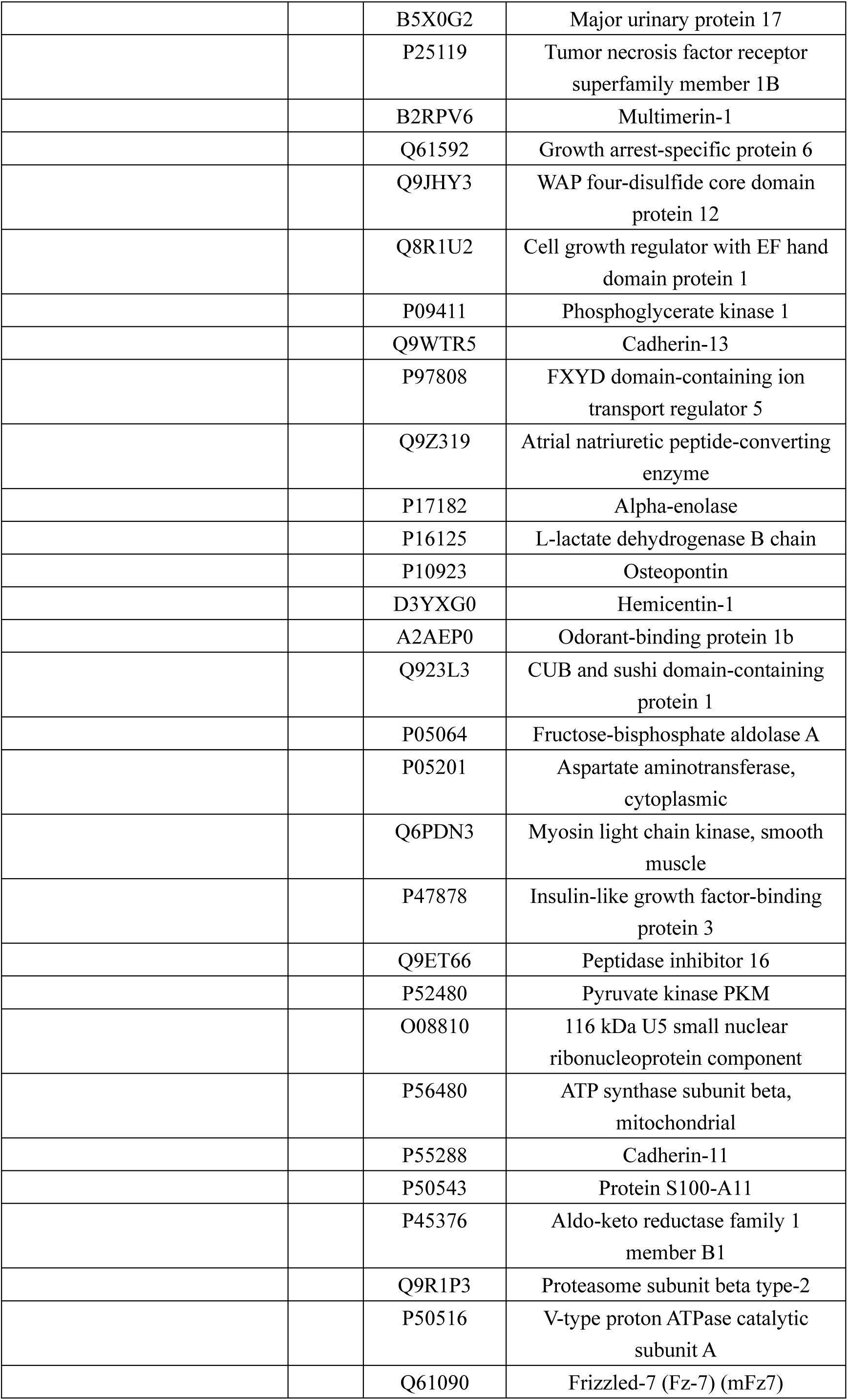
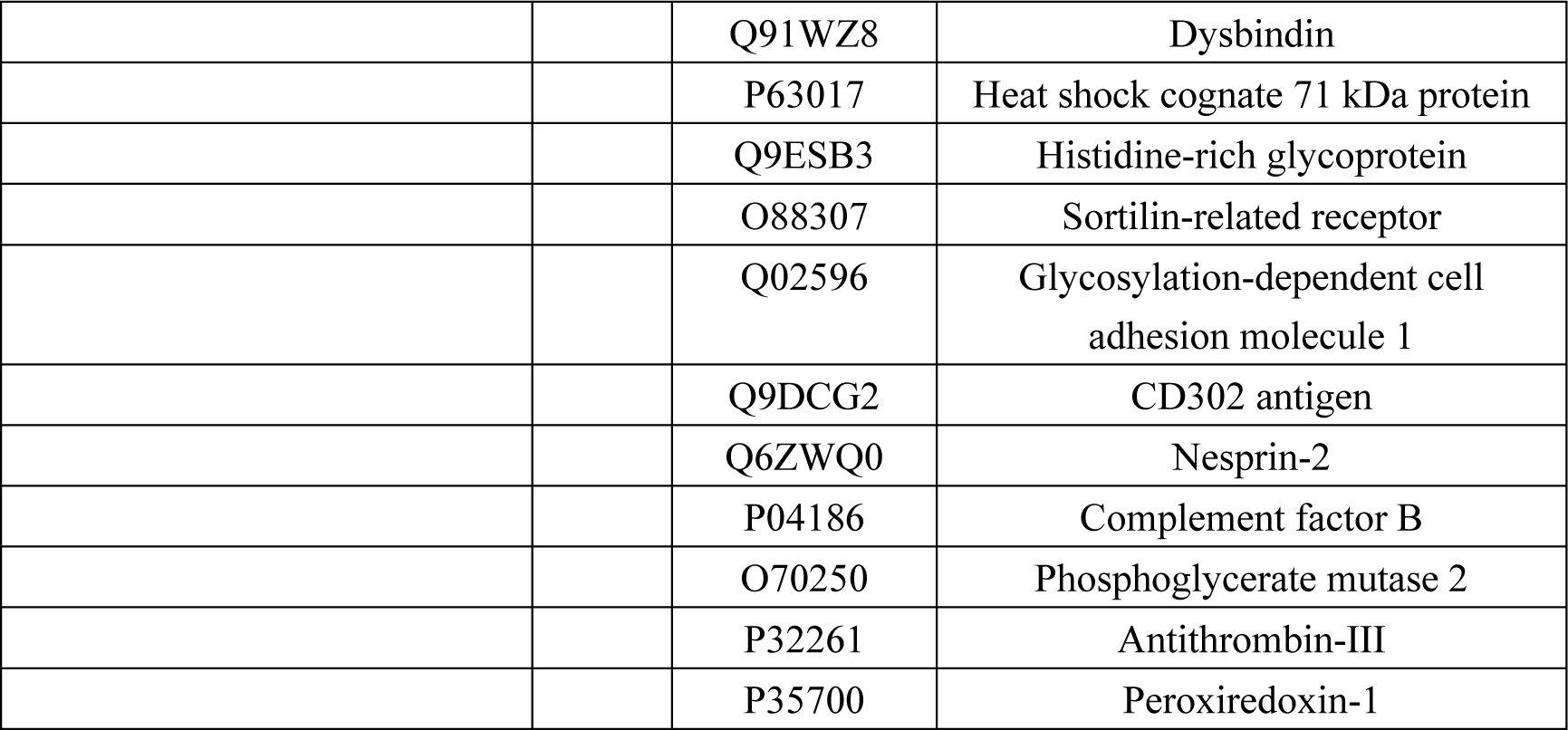
Shared differential proteins analysed in groups of four sweet taste substances.

**Table 2.** Shared biological processes analysed in groups of four sweet taste substances.

Exhibit 2 Shared biological processes analysed in groups of four sweet taste substances
| Names | total | elements |
| --- | --- | --- |
| acesulfame stevia glycosides<br>sucralose sucrose | 1 | positive regulation of cell migration |
| acesulfame sucralose sucrose | 3 | cell adhesion |
|  |  | blood coagulation |
|  |  | negative regulation of peptidase activity |
| acesulfame sucrose | 3 | positive regulation of inflammatory response |
|  |  | collagen fibril organisation |
|  |  | angiogenesis |
| sucralose sucrose | 19 | energy reserve metabolic process |
|  |  | mitochondrion morphogenesis |
|  |  | negative regulation of transcription, DNA-templated |
|  |  | cellular response to lipid |
|  |  | negative regulation of lipid storage |
|  |  | locomotor rhythm |
|  |  | response to bacterium |
|  |  | positive regulation of lipid metabolic process |
|  |  | positive regulation of MAPK cascade |
|  |  | negative regulation of lipid biosynthetic process |
|  |  | negative regulation of insulin secretion involved in cellular response to glucose stimulus |
|  |  | aerobic respiration |
|  |  | positive regulation of glucose metabolic process |
|  |  | negative regulation of gluconeogenesis |
|  |  | heat generation |
|  |  | Cellular response to testosterone stimulus |
|  |  | positive regulation of protein kinase B signaling |
|  |  | glucose homeostasis |
|  |  | positive regulation of gene expression |
| acesulfame stevia glycosides | 3 | peripheral nervous system axon regeneration |
|  |  | negative regulation of platelet-derived growth factor<br>receptor signaling pathway |
|  |  | positive regulation of cell proliferation |
| stevia glycosides sucralose | 3 | regulation of cell shape |
|  |  | mesenchyme migration |
|  |  | skeletal muscle thin filament assembly |
| acesulfame sucralose | 4 | gluconeogenesis |
|  |  | negative regulation of apoptotic process |
|  |  | carbohydrate metabolic process |
|  |  | positive regulation of ERK1 and ERK2 cascade |
| sucrose | 34 | platelet aggregation |
|  |  | regulation of cardiac conduction |
|  |  | response to cytokine |
|  |  | vascular smooth muscle cell differentiation |
|  |  | response to chromate |
|  |  | Regulation of systemic arterial blood pressure by renin-<br>angiotensin |
|  |  | angiotensin-activated signaling pathway |
|  |  | positive regulation of neuron projection development |
|  |  | Skin development |
|  |  | fibrinolysis |
|  |  | response to lead ion |
|  |  | response to estrogen |
|  |  | positive regulation of CoA-transferase activity |
|  |  | regulation of gene expression |
|  |  | ERK1 and ERK2 cascade |
|  |  | kidney development |
|  |  | plasminogen activation |
|  |  | angiotensin-mediated drinking behaviour |
|  |  | mesenchymal cell differentiation |
|  |  | skeletal system development |
|  |  | cell-matrix adhesion |
|  |  | extracellular matrix organisation |
|  |  | Learning or memory |
|  |  | one-carbon metabolic process |
|  |  | proteolysis |
|  |  | blood coagulation, fibrin clot formation |
|  |  | response to peptide hormone |
|  |  | response to methanol |
|  |  | positive regulation of epithelial to mesenchymal<br>transition |
|  |  | complement activation, alternative pathway |
|  |  | positive regulation of gap junction assembly |
|  |  | regulation of cell adhesion |
|  |  | negative regulation of endopeptidase activity |
|  |  | negative regulation of serine-type endopeptidase activity |
| stevia glycosides | 20 | regulation of centromere complex assembly |
|  |  | DNA-templated transcription, initiation |
|  |  | negative regulation of plasma membrane long-chain fatty acid transport |
|  |  | regulation of gene silencing |
|  |  | DNA replication-independent nucleosome assembly |
|  |  | cell proliferation |
|  |  | tetrahydrobiopterin biosynthetic process |
|  |  | nucleosome assembly |
|  |  | Wnt signaling pathway, calcium modulating pathway |
|  |  | DNA replication-dependent nucleosome assembly |
|  |  | Wnt signaling pathway |
|  |  | sensory perception of sound |
|  |  | actin filament network formation |
|  |  | actin filament bundle assembly |
|  |  | pericentric heterochromatin assembly |
|  |  | positive regulation of epithelial cell proliferation involved in wound healing |
|  |  | regulation of gene expression, epigenetic |
|  |  | negative regulation of chromosome condensation |
|  |  | negative regulation of megakaryocyte differentiation |
|  |  | L-phenylalanine metabolic process |
| acesulfame | 20 | Cellular response to reactive oxygen species |
|  |  | response to lipopolysaccharide |
|  |  | regulation of ruffle assembly |
|  |  | lipid metabolic process |
|  |  | positive regulation of cellular extravasation |
|  |  | positive regulation of NF-kappaB transcription factor activity |
|  |  | animal organ morphogenesis |
|  |  | metabolic process |
|  |  | positive regulation of phospholipase C activity |
|  |  | cell-cell adhesion |
|  |  | palate development |
|  |  | protein stabilisation |
|  |  | chondrocyte differentiation |
|  |  | skeletal system morphogenesis |
|  |  | positive regulation of peptidyl-tyrosine phosphorylation |
|  |  | microvillus assembly |
|  |  | positive regulation of nitric oxide biosynthetic process |
|  |  | positive regulation of kinase activity |
|  |  | protein localisation to plasma membrane |
|  |  | Homophilic cell adhesion via plasma membrane adhesion molecules |
| sucralose | 65 | histone H4 acetylation |
|  |  | positive regulation of binding |
|  |  | synaptic vesicle endocytosis |
|  |  | positive regulation of transcription, DNA-templated |
|  |  | negative regulation of cell migration |
|  |  | positive regulation of muscle contraction |
|  |  | striated muscle contraction |
|  |  | actin cytoskeleton organisation |
|  |  | morphogenesis of a polarised epithelium |
|  |  | establishment of Golgi localisation |
|  |  | canonical glycolysis |
|  |  | glycolytic process through fructose-6-phosphate |
|  |  | methylglyoxal biosynthetic process |
|  |  | positive regulation of protein phosphorylation |
|  |  | ATP biosynthetic process |
|  |  | retina development in camera-type eye |
|  |  | glycolytic process |
|  |  | cellular response to electrical stimulus |
|  |  | glyceraldehyde-3-phosphate biosynthetic process |
|  |  | histone H2A acetylation |
|  |  | axonogenesis |
|  |  | muscle cell cellular homeostasis |
|  |  | postsynaptic actin cytoskeleton organisation |
|  |  | extrinsic apoptotic signaling pathway |
|  |  | NAD metabolic process |
|  |  | positive regulation of T cell differentiation |
|  |  | regulation of protein localisation to plasma membrane |
|  |  | cellular phosphate ion homeostasis |
|  |  | regulation of blood coagulation |
|  |  | protein localisation to adherens junction |
|  |  | establishment or maintenance of cell polarity |
|  |  | cell motility |
|  |  | negative regulation of neuron death |
|  |  | positive regulation of cell motility |
|  |  | hemostasis |
|  |  | Cellular response to interleukin-7 |
|  |  | negative regulation of cyclin-dependent protein kinase activity |
|  |  | metanephric collecting duct development |
|  |  | adherens junction assembly |
|  |  | protein homotetramerisation |
|  |  | platelet activation |
|  |  | peripheral nervous system myelin maintenance |
|  |  | negative regulation of protein binding |
|  |  | cytolysis by host of symbiont cells |
|  |  | positive regulation of phosphorylation |
|  |  | ATP metabolic process |
|  |  | aging |
|  |  | positive regulation of double-strand break repair via homologous recombination |
|  |  | negative regulation of cell growth |
|  |  | negative regulation of hypoxia-induced intrinsic apoptotic signaling pathway |
|  |  | regulation of transmembrane transporter activity |
|  |  | Notch signaling pathway |
|  |  | regulation of multicellular organism growth |
|  |  | response to mechanical stimulus |
|  |  | regulation of norepinephrine uptake |
|  |  | Cellular response to cytochalasin B |
|  |  | cellular iron ion homeostasis |
|  |  | memory |
|  |  | positive regulation of plasminogen activation |
|  |  | protein trimerisation |
|  |  | positive regulation of phagocytosis |
|  |  | positive regulation of cytokine-mediated signaling pathway |
|  |  | Regulation of cyclin-dependent protein serine/threonine kinase activity |
|  |  | response to heat |
|  |  | apical protein localisation |

**Table 3.** Molecular functions of shared differential proteins analysed in groups of four sweet taste substances.

| Names | total | elements |
| --- | --- | --- |
| acesulfame<br>stevia glycosides<br>sucralose<br>sucrose | 1 | hydrolase activity |
| acesulfame<br>sucralose<br>sucrose | 4 | peptidase inhibitor activity |
|  |  | identical protein binding |
|  |  | receptor binding |
|  |  | serine-type endopeptidase inhibitor activity |
| acesulfame<br>stevia glycosides<br>sucralose | 1 | calcium ion binding |
| acesulfame<br>sucrose | 1 | endopeptidase inhibitor activity |
| sucralose<br>sucrose | 7 | peptidase activity |
|  |  | odourant binding |
|  |  | protease binding |
|  |  | small molecule binding |
|  |  | pheromone binding |
|  |  | extracellular matrix structural constituent |
|  |  | insulin-activated receptor activity |
| acesulfame<br>stevia glycosides | 2 | peroxiredoxin activity |
|  |  | thioredoxin peroxidase activity |
| stevia glycosides<br>sucralose | 1 | Wnt-protein binding |
| sucrose | 12 | endopeptidase activity |
|  |  | heparin binding |
|  |  | fibronectin binding |
|  |  | vitamin D binding |
|  |  | hydro-lyase activity |
|  |  | heparan sulfate proteoglycan binding |
|  |  | carbonate dehydratase activity |
|  |  | peptidase activator activity |
|  |  | arylesterase activity |
|  |  | proteoglycan binding |
|  |  | collagen binding |
|  |  | acetylcholine receptor binding |
| stevia glycosides | 7 | integrin binding |
|  |  | structural constituent of chromatin |
|  |  | chromatin binding |
|  |  | protein domain specific binding |
|  |  | PDZ domain binding |
|  |  | DNA binding |
|  |  | protein heterodimerisation activity |
| acesulfame | 15 | growth factor binding |
|  |  | macromolecular complex binding |
|  |  | carbohydrate binding |
|  |  | protein binding involved in protein folding |
|  |  | beta-galactosidase activity |
|  |  | signaling receptor activity |
|  |  | hydrolase activity, acting on glycosyl bonds |
|  |  | protein tyrosine kinase activity |
|  |  | beta-amyloid binding |
|  |  | syndecan binding |
|  |  | virus receptor activity |
|  |  | hydrolase activity, hydrolyzing O-glycosyl compounds |
|  |  | antioxidant activity |
|  |  | transmembrane receptor protein tyrosine kinase activity |
|  |  | protein homodimerisation activity |
| sucralose | 23 | ADP binding |
|  |  | intramolecular transferase activity, phosphotransferases |
|  |  | serine-type peptidase activity |
|  |  | Structural constituent of postsynaptic actin cytoskeleton |
|  |  | enzyme binding |
|  |  | cadherin binding |
|  |  | catalytic activity |
|  |  | nitric-oxide synthase binding |
|  |  | 2,3-bisphosphoglycerate-dependent phosphoglycerate mutase activity |
|  |  | nucleotide binding |
|  |  | Tat protein binding |
|  |  | heat shock protein binding |
|  |  | ATP binding |
|  |  | Wnt-activated receptor activity |
|  |  | fructose binding |
|  |  | fructose-bisphosphate aldolase activity |
|  |  | bisphosphoglycerate mutase activity |
|  |  | phosphopyruvate hydratase activity |
|  |  | growth factor receptor binding |
|  |  | chaperone binding |
|  |  | kinesin binding |
|  |  | protein binding |
|  |  | phosphoglycerate mutase activity |

**Table 4.** Shared differential proteins common to individual analyses of the four sweet substances.

| Names | total | Uniprot ID | Protein name |
| --- | --- | --- | --- |
| acesulfame<br>stevia glycosides<br>sucralose | 1 | Q60928 | Glutathione hydrolase 1 proenzyme |
| acesulfame stevia<br>glycosides | 1 | P52760 | 2-iminobutanoate/2-iminopropanoate deaminase |
| acesulfame<br>sucralose | 2 | P70441 | Na(+)/H(+) exchange regulatory cofactor NHE-RF1 |
|  |  | P12658 | Calbindin |
| sucrose | 1 | Q61391 | Enkephalinase |
| acesulfame | 10 | Q91ZI0 | Cadherin EGF LAG seven-pass G-type receptor 3 |
|  |  | Q9JI02 | Secretoglobin family 2B member 20 |
|  |  | Q60932 | Voltage-dependent anion-selective channel protein<br>1 |
|  |  | P99029 | Peroxiredoxin-5, mitochondrial |
|  |  | Q9DAU7 | WAP four-disulfide core domain protein 2 |
|  |  | Q91XE4 | N-acyl-aromatic-L-amino acid amidohydrolase |
|  |  | P23780 | Beta-galactosidase |
|  |  | P70699 | Lysosomal alpha-glucosidase |
|  |  | Q06318 | Uteroglobin |
|  |  | Q9JHY4 | WAP four-disulfide core domain protein 15B |
| sucralose | 8 | P11087 | Collagen alpha-1(I) chain |
|  |  | P04939 | Major urinary protein 3 |
|  |  | P15626 | Glutathione S-transferase Mu 2 |
|  |  | P05784 | Keratin, type I cytoskeletal 18 |
|  |  | Q02596 | Glycosylation-dependent cell adhesion molecule 1 |
|  |  | P41272 | CD27 antigen |
|  |  | P10923 | Osteopontin |
|  |  | P09411 | Phosphoglycerate kinase 1 |

## Reference

1 Walton J, Bell H, Re R, Nugent AP. Current perspectives on global sugar consumption: definitions, recommendations, population intakes, challenges Nutr Res Rev. 2023;36(1):1–22.

2 Lenoir M, Serre F, Cantin L, Ahmed SH. Intense sweetness surpasses cocaine reward. PLoS One. 2007;2(8):e698.

3 Clara R. Freeman, Amna Zehra, Veronica Ramirez, Corinde E. Wiers, Nora D. Volkow, Gene-Jack Wang. impact of sugar on the body, brain, and behaviour. front. biosci. 2018, 23(12), 2255–2266.

4 Ruanpeng D, Thongprayoon C, Cheungpasitporn W, Harindhanavudhi T. Sugar and artificially sweetened beverages linked to obesity: a systematic review and meta-analysis. qjm. 2017;110(8):513–520.

5 Imamura F, O’Connor L, Ye Z, et al. Consumption of sugar sweetened beverages, artificially sweetened beverages, and fruit juice and incidence of type 2 diabetes: systematic review, meta-analysis, and estimation of population attributable fraction. bmj. 2015;351:h3576.

6 Xi B, Huang Y, Reilly KH, et al. Sugar-sweetened beverages and risk of hypertension and CVD: a dose-response meta-analysis. Br J Nutr. 2015;113(5):709–717.

7 Asgari-Taee F, Zerafati-Shoae N, Dehghani M, Sadeghi M, Baradaran HR, Jazayeri S. Association of sugar sweetened beverages consumption with non alcoholic fatty liver disease: a systematic review and meta-analysis. Eur J Nutr. 2019;58(5):1759–1769.

8 Valenzuela MJ, Waterhouse B, Aggarwal VR, Bloor K, Doran T. Effect of sugar-sweetened beverages on oral health: a systematic review and meta-analysis. Eur J Public Health. 2021;31(1):122–129.

9 Llaha F, Gil-Lespinard M, Unal P, de Villasante I, Castañeda J, Zamora-Ros R. Consumption of Sweet Beverages and Cancer Risk. a Systematic Review and Meta-Analysis of Observational Studies. nutrients. 2021;13(2):516.

10 Lange FT, Scheurer M, Brauch HJ. Artificial sweeteners--a recently recognised class of emerging environmental contaminants: a review. Anal Bioanal Chem. 2012;403(9):2503–2518.

11 Praveena SM, Cheema MS, Guo HR. Non-nutritive artificial sweeteners as an emerging contaminant in the environment: a global review and risks perspectives. Ecotoxicol Environ Saf. 2019;170:699–707.

12 Fung TT, Malik V, Rexrode KM, Manson JE, Willett WC, Hu FB. Sweetened beverage consumption and risk of coronary heart disease in women. Am J Clin Nutr. 2009;89:1037–42.

13 Lin J, Curhan GC. Associations of sugar and artificially sweetened soda with albuminuria and kidney function decline in women. Clin J Am Soc Nephrol. 2011;6:160–6.

14 Pepino MY, Tiemann CD, Patterson BW, Wice BM, Klein S. Sucralose affects glycemic and hormonal responses to an oral glucose load. Diabetes Care. 2013. 36(9):2530–2535.

15 Gao, Youhe. Does urine have the potential to be a biomarker goldmine? Sci. China Life Sci. 2013, 43(8): 708–708

16 Logue C, Dowey LC, Strain JJ, Verhagen H, Gallagher AM. The potential application of a biomarker approach for the investigation of low-calorie sweetener exposure. Proceedings of the Nutrition Society. 2016;75(2):216–225.

17 Sclafani A, Bahrani M, Zukerman S, Ackroff K. Stevia and saccharin preferences in rats and mice. Chem Senses. 2010 Jun;35(5):433–43.

18 Yin KJ, Xie DY, Zhao L, Fan G, Ren JN, Zhang LL, Pan SY. Effects of different sweeteners on behaviour and neurotransmitters release in mice. J Food Sci Technol. 2020 Jan;57(1):113–121.

19 Gardner EL. Addiction and brain reward and antireward pathways. Adv Psychosom Med. 2011;30:22–60.

